# Targeted RNA Condensation in Living Cells via Genetically Encodable Triplet Repeat Tags

**DOI:** 10.1101/2023.04.07.536084

**Authors:** Zhaolin Xue, Kewei Ren, Rigumula Wu, Zhining Sun, Ru Zheng, Qian Tian, Ahsan Ausaf Ali, Lan Mi, Mingxu You

## Abstract

Living systems contain various functional membraneless organelles that can segregate selective proteins and RNAs via liquid–liquid phase separation. Inspired by nature, many synthetic compartments have been engineered in vitro and in living cells, mostly focused on protein-scaffolded systems. Herein, we introduce a nature-inspired genetically encoded RNA tag to program cellular condensate formations and recruit non-phase-transition target RNAs to achieve functional modulation. In our system, different lengths of CAG-repeat tags were tested as the self-assembled scaffold to drive multivalent condensate formation. Various selective target messenger RNAs and noncoding RNAs can be compartmentalized into these condensates. With the help of fluorogenic RNA aptamers, we have systematically studied the formation dynamics, spatial distributions, sizes, and densities of these cellular RNA condensates. The regulation functions of these CAG-repeat tags on the cellular RNA localization, lifetime, RNA–protein interactions, and gene expression have also been investigated. Considering the importance of RNA condensation in both health and disease conditions, these genetically encodable modular and self-assembled tags can be potentially widely used for chemical biology and synthetic biology studies.

## Introduction

The functions of cellular RNAs are highly related with their subcellular localizations and local environment. One ubiquitous approach to control RNA localization is via macromolecular condensation, i.e., the formation of membraneless subcellular compartments. RNA compartmentalization is prevalent and plays critical roles in process such as transcription, splicing, RNA degradation, heterochromatin formation, and stress response.^1–4^ The precise modulation of subcellular compartmentalization of specific RNA sequences is thus important for controlling gene expression and cellular functions.

Cellular RNA localization, compartmentalization, and trafficking have been largely regulated by the formation of ribonucleoprotein (RNP) complexes via RNA–protein interactions.^5,6^ Recent studies demonstrated that specific RNA self-assemblies, particularly among repeat expansions, can also be used to mediate RNA phase separation inside living cells.^7–10^ Compared to protein-based RNA compartmentalization, RNA–RNA interaction-mediated condensate formation can be highly sequence-specific, modular, and programmable.^11,12^ As a result, powerful RNA devices may be engineered to control cellular compartmentalization of specific cellular RNAs and modulate their local concentrations and functions.

In this study, short CAG trinucleotide repeats are engineered into genetically encodable self-regulated tags to recruit and condense different cellular target RNAs. CAG repeats are used here because these trinucleotides can readily form condensation, without the involvement of proteins.^9,10,13,14^ We want to test here if these naturally occurring CAG repeats can be used as functional RNA nanodevices, either being a cis-acting RNA element within target RNA transcript or as a trans-acting effector functioning through specific hybridization with target RNAs. Both cis- and trans-mechanisms can be potentially applied to develop general platforms to induce the phase separation of various target RNAs, in vitro and in living systems.

To image these RNA condensates, especially inside living cells, we fuse the CAG-repeat-tagged RNA strands with a fluorogenic RNA aptamer, named Broccoli. Broccoli is an RNA strand that can selectively bind and activate the fluorescence signal of small-molecule dyes, such as DFHBI-1T.^15^ The dynamics, sizes, and densities of RNA condensates can thus be visualized and quantified based on fluorescence images. Our results show that target RNAs of different lengths and sequences can be efficiently recruited into condensates by these CAG-repeat tags. CAG repeats can also be genetically encoded to regulate the subcellular localization, compartmentalization, and function of various messenger RNAs (mRNAs) and non-coding RNAs inside living bacterial cells.

## Results and Discussion

We first asked if Broccoli can be used to visualize CAG-repeat-induced RNA condensation. To test this, we synthesized F30-2d×Broccoli-tagged RNA strands containing 0×, 4×, 20×, 31×, or 47×CAG repeats, which were named as **0R**, **4R**, **20R**, **31R**, and **47R**, respectively. The F30 scaffold was used to ensure the proper folding of two incorporated dimeric Broccoli RNA (Table S1).^16^ Compared to untagged F30-2d×Broccoli (i.e., 0R), after attaching these RNA repeats, F30-2d×Broccoli still exhibits strong, even slightly higher, fluorescence signals (Figure S1a). By annealing a solution containing 4 μM RNA, 20 mM MgCl_2_, and 80 μM DFHBI-1T, 20R, 31R, and 47R samples exhibited obvious RNA condensation (Figure 1a), with a large number of spherical-shaped fluorescent condensates at ~1.4–1.8 μm in diameter (Figure 1c). In contrast, almost no phase separation was observed with the 0R and 70AC control. Here, **70AC** is an F30-2d×Broccoli-tagged RNA of equivalent length as 47R but contains only repeated AC dinucleotides. Interestingly under our experimental condition, with only 4×CAG repeats, 4R can also generate large-sized RNA condensates (diameter, ~1.8 μm), while at a lower density than longer CAG repeats.

**Figure 1.**
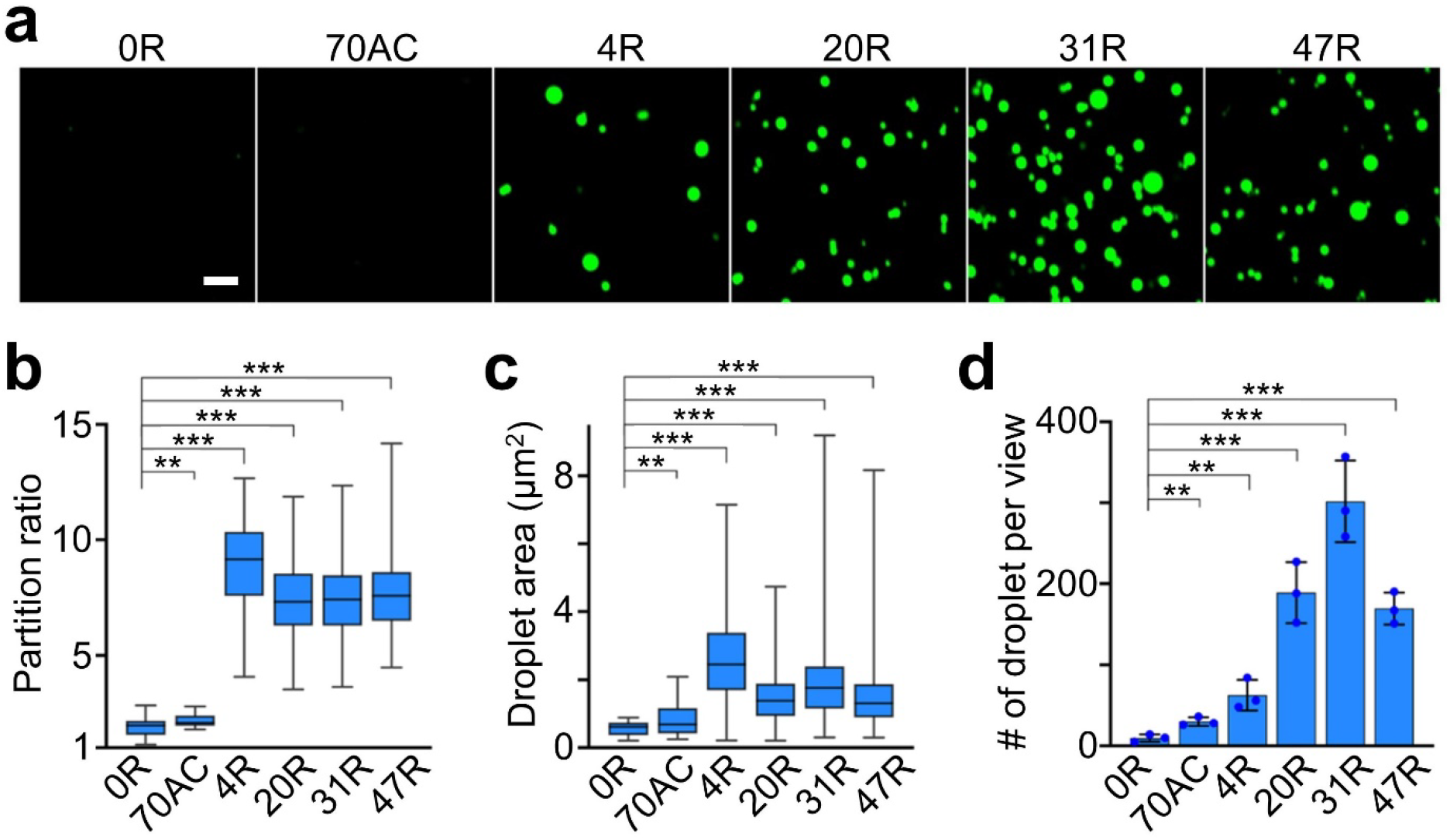
CAG repeats-regulated *in vitro* RNA condensation. (**a**) Confocal fluorescence imaging of RNA condensates induced by F30-2d×Broccoli-tagged 4×, 20×, 31×, or 47×CAG repeats (4R, 20R, 31R or 47R). F30-2d×Broccoli (0R) and F30-2d×Broccoli-tagged 70 repeats of AC (70AC) were used as the negative control. Fluorescence images were taken in solutions containing 4 μM RNA, 20 mM MgCl_2_, and 80 μM DFHBI-1T after annealing. Scale bar, 5 μm. (**b**) The partition ratio, i.e., the ratio of average fluorescence intensity inside individual condensate versus background solution signals, and (**c**) droplet area of each type of RNA condensate is plotted in box with min-to-max whiskers. A total of 20, 29, 241, 318, 323, and 437 condensates were analyzed in the case of 0R, 70AC, 4R, 20R, 31R, and 47R, respectively. (**d**) The number of droplets observed per imaging view. Each imaging view equals to 4,430 μm^2^. Shown are the mean and the standard deviation (SD) values from at least three representative images. Two-tailed student’s t-test: ***, *p*<0.001; **, *p*<0.01.

To further compare the RNA densities within these different CAG-repeat aggregates, we characterized the partition ratio of each single condensate, which is defined as the ratio of average fluorescence intensity inside individual condensate versus background solution fluorescence signals. A partition ratio of ~2.2, 8.9, 7.4, 7.3 and 7.7 was exhibited for the 70AC, 4R, 20R, 31R, and 47R condensates, respectively (Figure 1d). 4R exhibited a slightly higher partition ratio and larger aggregation size as compared to other expanded CAG repeats. On the other hand, 4R solution contained less condensates (~5 per 400 μm^2^ imaging area) than that of 20R, 31R, and 47R (>15 counts per same area) (Figure 1). These results indicated that the CAG repeats can indeed induce *in vitro* RNA phase separation. Meanwhile, Broccoli can be used as a fluorescence imaging reporter for RNA condensates, and these fluorescence signals may also be used to estimate RNA concentrations within each condensate (Figure S1b, c).

We noticed that these CAG-repeat condensates tend to exhibit gel-like behaviors, such as stacking and slow fusion (Figure 1a). These gel-like properties can be resulted from strong multivalence interactions among these trinucleotide repeats.^8,17^ A fluorescence recovery after photobleaching (FRAP) approach was also applied to measure RNA mobilities in condensates. Minimal fluorescence recovery was observed (over a total of ~5 min, Figure S2), indicating that *in vitro* formed CAG condensates indeed display a highly static structure. The kinetics of RNA condensation was also monitored right after the snap cooling of a solution containing 4 μM 47R and 20 mM MgCl_2_. Our results showed that the number of RNA condensates kept increasing during the first 15 min of incubation, while the diameter and partition ratio of each condensate already reached ~90% of the maximum level in ~5 min (Figure S3). These data indicated the fast assembly kinetics of these RNA condensates.

We also studied the effect of RNA and Mg^2+^ concentrations on the CAG-repeat-induced phase separation. In our test, 0.02–10 μM of 0R, 20R, 31R, or 47R strand was separately mixed with 5–40 mM of MgCl_2_. As expected, the formation of RNA condensates can be facilitated with increasing concentrations of RNA and MgCl_2_ (Figure S4). RNAs with longer CAG repeats, e.g., 31R and 47R, can more readily form condensates, even at reduced RNA and MgCl_2_ levels. It is worth mentioning that in these tests, almost no RNA condensation was observed under physiologically relevant ≤5 mM Mg^2+^ ion conditions.^18,19^ Meanwhile, minimal RNA condensation was shown without annealing. However, intracellular RNA compartmentalization can still be quite different from these in vitro tests, as RNA condensation can be potentially facilitated by the crowded and protein-rich cellular environment.^7,20,21^

Before testing RNA condensation in living cells, we also wondered if these CAG repeats can function as general molecular tags to induce the phase transition of attached RNA sequences. For this purpose, we synthesized 20 different RNA strands that contain 5’-F30-2d×Broccoli-tagged 100-, 200-, 500-, 1,000-, or 2,000-nucleotide (nt)-long scrambled sequences. 0×, 20×, 31×, or 47×CAG repeats were conjugated at the 3’-end of these strands, respectively. 4×CAG repeats were not chosen because their condensation occurs at low efficiency. Without attaching CAG repeats, scrambled RNAs (named as **0.1k**, **0.2k**, **0.5k**, **1.0k**, and **2.0k**) could not form condensates in a solution containing 4 μM RNA, 20 mM MgCl_2_, and 80 μM DFHBI-1T (Figure 2). In contrast, the 20×CAG tag can induce compartmentalization when short RNA sequences (i.e., 100 and 200 nt) were attached, while obvious condensates were shown in all the solutions containing 31×CAG- or 47×CAG-tagged RNA strands. These results indicated that CAG repeats can still induce phase separation even after tagging with long non-condensation RNA strands.

**Figure 2.**
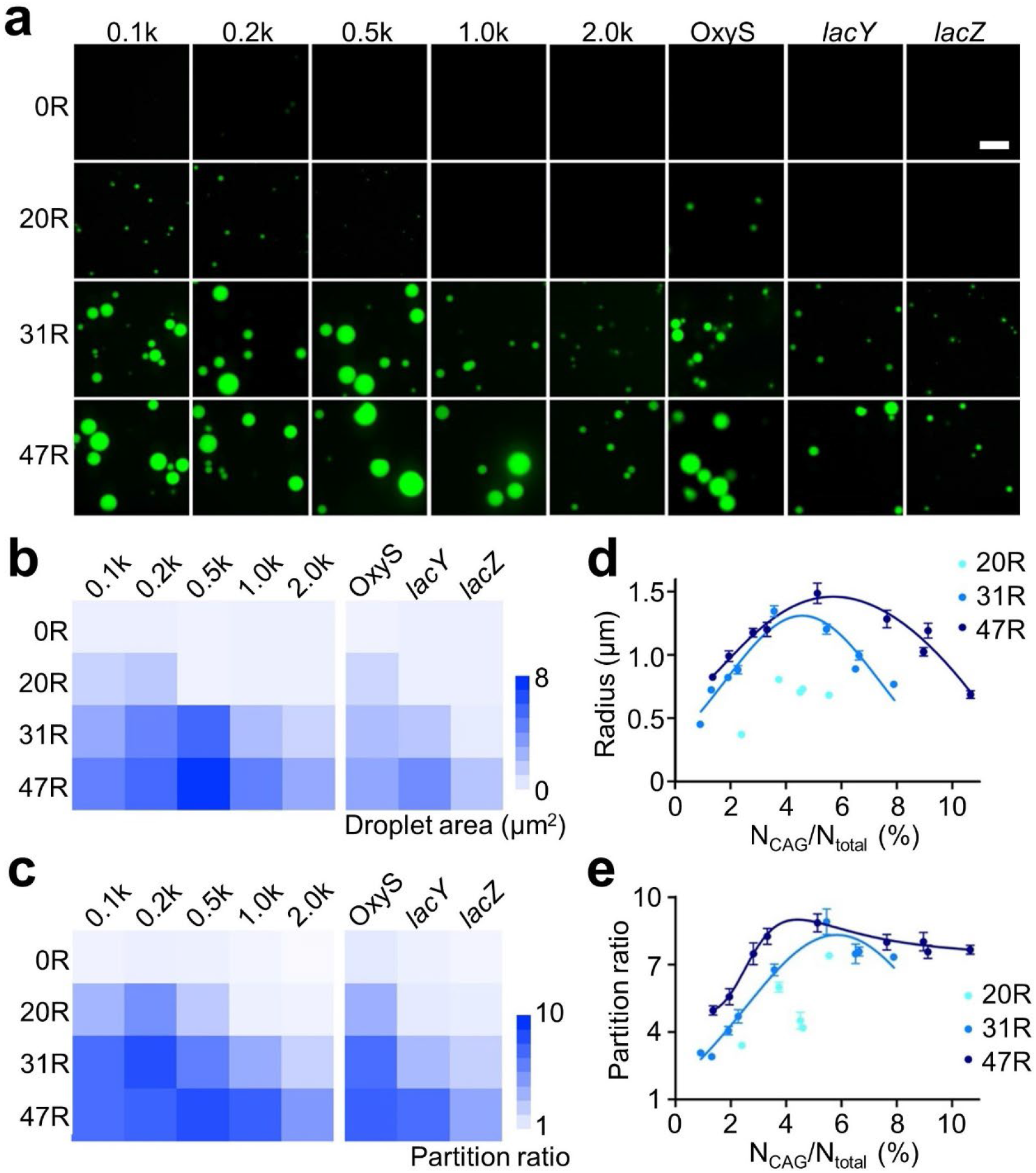
Targeted RNA condensation via CAG repeats. (**a**) Confocal fluorescence imaging of F30-2d×Broccoli-0×, 20×, 31×, or 47×CAG repeats (0R, 20R, 31R, 47R)-tagged 100-, 200-, 500-, 1,000-, or 2,000-nt-long scrambled RNA (0.1k, 0.2k, 0.5k, 1.0k, 2.0k), or OxyS (109 nt), *lacY* (1,254 nt), *lacZ* (3,044 nt) RNA sequences. Fluorescence images were taken in solutions containing 4 μM RNA, 20 mM MgCl_2_, and 80 μM DFHBI-1T after annealing. Scale bar, 5 μm. (**b**) The mean droplet area and (**c**) partition ratio of individual condensate is plotted based on data shown in Figure S5 and S6. (**d**) The radius and (**e**) partition ratio of each studied condensate is plotted as a function of the ratio between the number of CAG repeats and total RNA length. Results from each 20×, 31× or 47×CAG-tagged target RNA, including F30-2d×Broccoli, 0.1k, 0.2k, 0.5k, 1.0k, 2.0k scrambled RNAs, and OxyS, *lacY* and *lacZ* are gathered and denoted under 20R (cyan), 31R (light blue) and 47R (dark blue), respectively. Shown are the mean and the standard error of the mean (SEM) values from at least three replicated measurements for each data point.

We further quantified the correlations between the length of CAG repeats and the partition ratio and size of condensates. As shown in Figure S5, a longer CAG-repeat tag can generally increase both the diameter and partition ratio of RNA condensates. While interestingly, after attaching longer scrambled RNAs to the same CAG-repeat tag, the partition ratio and size of condensates tended to first increase and then decrease (Figure 2b, c). This result is consistent with those observed in synthetic multivalent polymers with changing valency of interactions,^22^ suggesting that the efficiency of condensate formation is influenced by the sequence length of both CAG repeats and target RNAs. A longer CAG repeat tag is normally needed to recruit larger RNA targets into condensates.

To next study if CAG repeats can also mediate condensation of endogenous RNA sequences, three bacterial RNAs were tested, including a 109-nt-long OxyS small noncoding RNA (sRNA) and two *lac* operon mRNA, *lacY* (1,254 nt) and *lacZ* (3,044 nt). After tagging with F30-2d×Broccoli and 20×, 31×, or 47×CAG repeats, OxyS can easily form obvious condensates, with size and partition ratio comparable to those of 0.1k scrambled RNAs (Figure 2 and S6). Similarly, 31× and 47×CAG-tagged *lacY* mRNA can also generate condensates close to those of 1.0k scrambled RNAs. Even in the case of the 3,044-nt-long *lacZ* mRNA, after conjugating with 31×CAG and 47×CAG, micrometer-sized condensates can be clearly observed (Figure 2a). These data demonstrated that the CAG repeat tags can facilitate both scrambled and functional RNA species to partition into condensates.

By combining all these results from different RNA targets, our data showed that the size of condensates tended to be first enlarged and then reduced with an increasing ratio between the CAG repeat number and total RNA length, **NCAG/Ntotal** (Figure 2d). Meanwhile, the partition ratio is also generally increased at low NCAG/Ntotal ratio and then slightly decreased at NCAG/Ntotal >5% for 31× and 47×CAG repeat samples (Figure 2e). These results suggested that NCAG/Ntotal ratio can be potentially an important factor for regulating the aggregation status of RNA-repeat condensates.

After all these *in vitro* characterizations, we next asked if CAG repeats could also be used as genetically encoded tags to regulate the condensation of target RNAs inside living cells. In our test, we first transformed BL21 Star^TM^ (DE3) *E. coli* cells with pET-28c vectors that express 4R, 20R, 31R, or 47R sequences. Compared to the control cells encoding only F30-2d×Broccoli (0R) or 70AC, the formation of cellular condensates can be clearly visualized in CAG-repeat-containing strains (Figure 3a). Similar to the *in vitro* results, the number and partition ratio of cellular condensates highly depend on the length of CAG repeats. Most abundant and RNA-concentrated condensates were shown in 47R-expressing cells (Figure 3b, c). Vast majority of CAG-repeat-expressing cells (4R–47R) contain one or two condensates (76%–85%), mainly localized at the cell poles. Meanwhile, it is worth mentioning that minimal cytotoxicity was observed in these CAG repeats-expressing *E. coli* cells (Figure S7). Indeed, CAG repeat-regulated formation of RNA condensates can occur in living bacterial cells.

**Figure 3.**
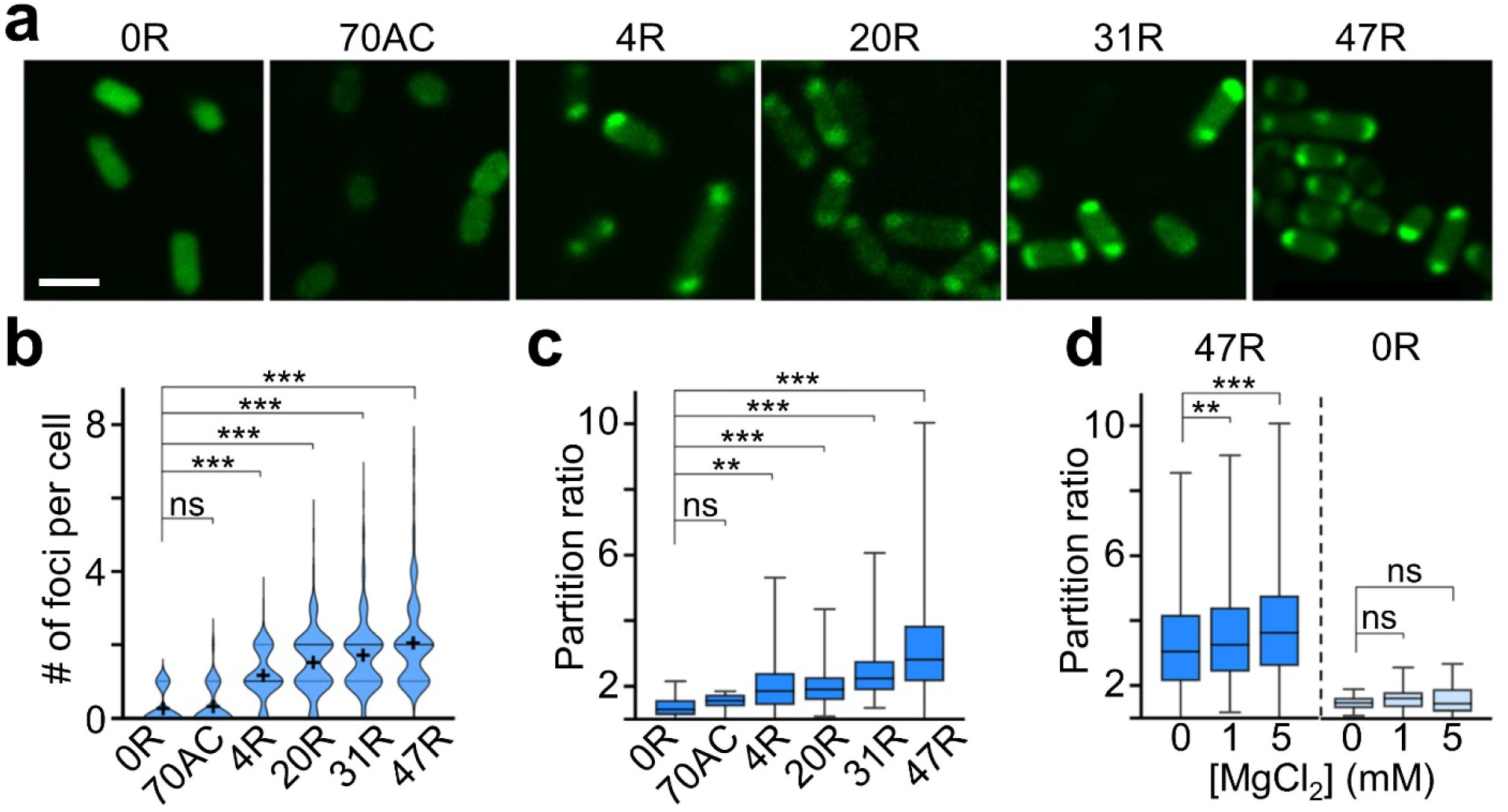
RNA condensate formation in living bacterial cells. (**a**) Confocal fluorescence imaging of BL21 Star^TM^ (DE3) *E. coli* cells that express F30-2d×Broccoli-tagged 0×, 4×, 20×, 31×, 47×CAG repeats (0R, 4R, 20R, 31R, 47R) or 70×AC repeats (70AC). Scale bar, 2 μm. (**b**) The violin plot distribution of the number of foci per cell. Solid and dashed line indicates the median and interquartile value, respectively. The black cross indicates the mean value. (**c**) The partition ratio of individual cellular foci as measured in DPBS buffer without adding MgCl_2_. A total of 20, 29, 241, 318, 323, and 437 condensates were analyzed in 0R, 70AC, 4R, 20R, 31R, and 47R cells, respectively. (**d**) Magnesium ion-dependent partition ratio changes of individual cellular foci. A total of 421, 562, 1058 (17, 21, 27) condensates in 47R (0R) cells were analyzed in the presence of 0, 1, or 5 mM Mg^2+^ respectively. Shown are box plots with min-to-max whiskers. All the data is collected from at least three representative images. Two-tailed student’s t-test: ***, *p*<0.001; **, *p*<0.01; ns, not significant, *p*>0.05.

We also applied FRAP to study the mobility of RNAs within these cellular condensates. In contrast to our *in vitro* data (Figure S2), these intracellular RNA condensates exhibit more liquid-like properties as fast fluorescence recovery was observed: ~90% of original fluorescence signals being reached in ~30 s (Figure S8). Interestingly, for cells possessing two major condensates at opposite poles, after photobleaching the condensate at one pole, a clear transfer of fluorescence signal from the other unbleached pole was observed in 5 out of 9 tested cells (Figure S8).

To further study if the formation of these cellular condensates can be simply regulated by adding magnesium ions to increase the binding affinities among CAG-repeat strands, we incubated 0R- and 47R-expressing cells with 0, 1, or 5 mM MgCl_2_. Indeed, both the number and partition ratio of RNA condensates were upregulated in 47R cells after adding 5 mM MgCl_2_ (Figure 3d and S9). In contrast, no changes were observed in 0R cells, meanwhile, the average cellular Broccoli fluorescence in both groups of cells was not altered. Mg^2+^ can thus be used as a convenient regulator of these cellular CAG-repeat condensates.

In addition, our data showed that the formation of CAG condensates may reduce the cellular RNA degradation in these bacterial cells. In our test, after incubation for 24 hours, ~45% of 47R cellular fluorescence could still be observed, with most signals coming from condensates. While in 0R-expressing *E. coli* cells, the cellular fluorescence was decreased by >75% under this same experimental condition (Figure S10).

We next tested if CAG repeats can also recruit other cellular RNA targets into condensates. For this purpose, we prepared pET-28c vectors that respectively express 0R, 20R, 31R, or 47R-tagged OxyS, *lacY,* and *lacZ* RNAs. After transforming into BL21 Star™ (DE3) cells, bright fluorescent foci can be observed only in CAG-repeat-expressing cells (Figure S11). Both the number and partition ratio of RNA condensates tend to be increased after attaching elongated CAG repeats (Figure S11). On average, ~1.5–2.4 condensates were shown in each individual cell, mostly at the poles. All these data supported that the CAG repeats can be used as genetically encoded tags to drive the phase transition of target RNAs inside living cells.

To further assess if the formation of CAG condensates will change the cellular localization of target RNAs, we applied super-resolution structured illumination microscopy to image 0R- and 47R-tagged *lacY* mRNA. It is known that *lacY* prefers to localize near bacterial membranes, i.e., the functioning site of its protein product, lactose permease LacY.^23–25^ Indeed, without the CAG-repeat modification, *lacY-0R* was exclusively located at the *E. coli* membranes, exhibiting a hollow fluorescent pattern around the cells (Figure 4a). In contrast, after tagging with 47R, the hollow membrane fluorescent pattern was disrupted and replaced by condensation across the cytoplasm in most majority of cells. These results further validated that the CAG-repeat tags can alter the cellular locations of the attached RNAs.

**Figure 4.**
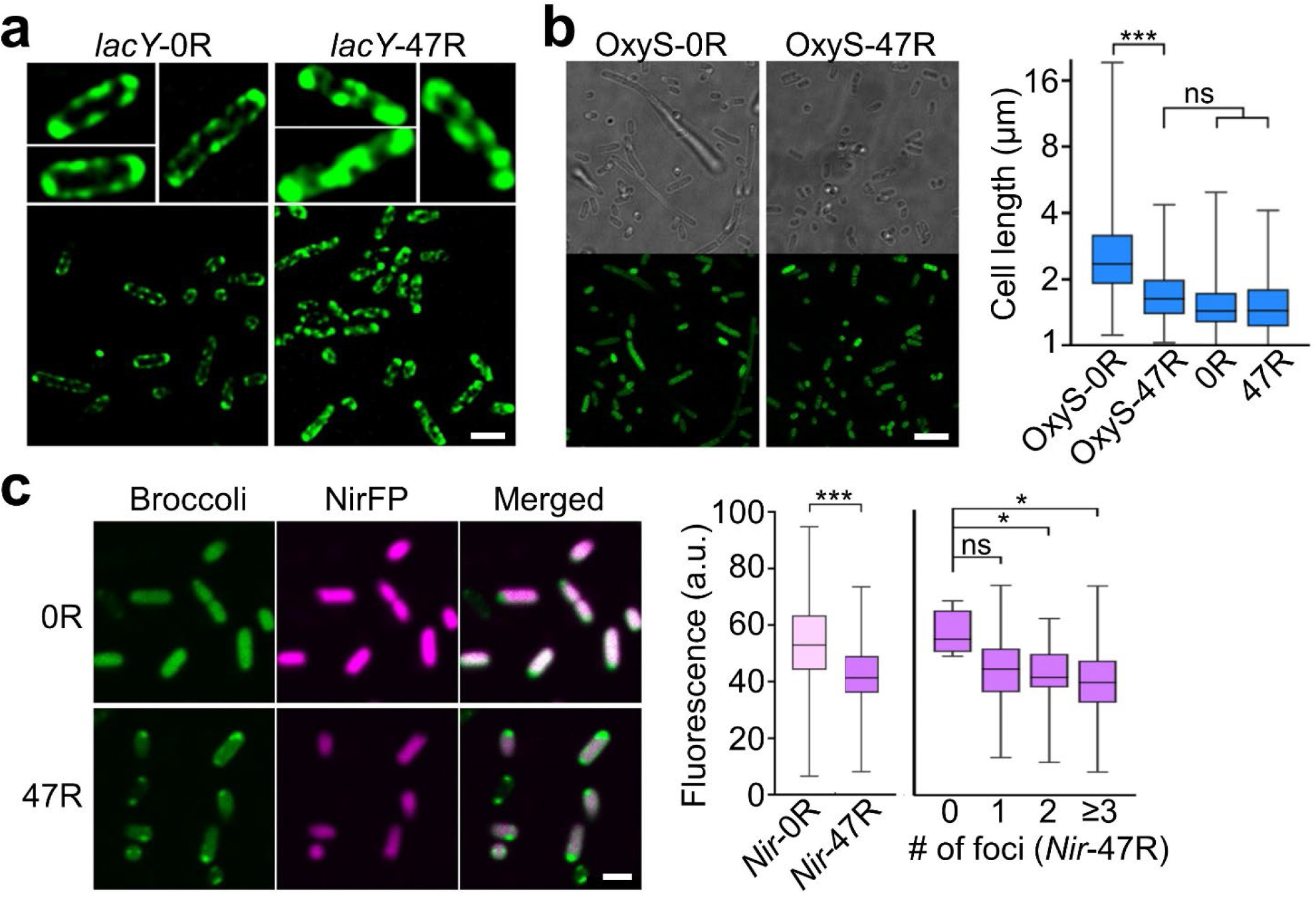
Condensation-regulated cellular localization and function of target RNAs. (**a**) Structured illumination microscope imaging of *E. coli* cells expressing 0R- or 47R-tagged *lacY* mRNA. Scale bar, 2 μm. (**b**) Confocal fluorescence imaging of *E. coli* cells expressing 0R- or 47R-tagged OxyS sRNA. Scale bar, 5 μm. Individual cell length is also plotted and compared to 0R- and 47R-expressing cells that without attaching OxyS. Shown are box plots with min-to-max whiskers. A total of 200, 153, 221, and 201 cells were analyzed in OxyS-0R, OxyS-47R, 0R, and 47R, respectively. (**c**) Confocal fluorescence imaging of *E. coli* cells expressing 0R- or 47R-tagged near-infrared fluorescent protein (NirFP) mRNA. Scale bar, 2 μm. Average cellular NirFP fluorescence as measured in 120 and 99 individual 0R- or 47R-tagged cells are plotted. Meanwhile, the NirFP fluorescence signals in individual 47R-tagged cells are also separately plotted based on the corresponding number of cellular foci. A total of 4, 25, 30, and 40 cells were analyzed with 0, 1, 2, and ≥3 foci, respectively. Shown are box plots with min-to-max whiskers. All the data is collected from at least three representative images. Two-tailed student’s t-test: ***, *p*<0.001; *, *p*<0.05; ns, not significant, *p*>0.05.

We also studied if the phase transition of target RNAs can be used to control their cellular functions. OxyS sRNA is known to repress the expression of transcription termination factor NusG and impair cell division, and as a result, generate long bacterial cells.^26,27^ By comparing the length of individual *E. coli* cells expressing either 0×CAG- or 47×CAG-tagged OxyS, our results showed that the average length of OxyS-47R cells is ~46% shorter than OxyS-0R cells (Figure 4b). As a control, without attaching OxyS, 0R- and 47R-expressing *E. coli* cells exhibited almost identical lengths as those of OxyS-47R cells. These data indicated that after condensation, the regulation function of OxyS on cell division is impaired, likely due to a reduced chance of OxyS to interact with *nusG* mRNA, which resumed the NusG expression.^26^

To further test if protein synthesis can indeed be regulated via the condensation of specific mRNAs, we transformed BL21 Star^TM^ (DE3) cells with vectors expressing 0×CAG- or 47×CAG-tagged 705-nt-long mRNA (named as ***Nir-0R*** and ***Nir*-47R**) that encodes a near-infrared fluorescent protein (NirFP). Compared to *Nir-0R* cells, ~20% lower NirFP signals were observed in *Nir*-47R cells (Figure 4c). Most *Nir*-47R RNA was located within condensates at the poles (as shown in the Broccoli channel), while the majority of translated NirFP proteins were observed throughout the cells, except the poles. We further plotted the cellular NirFP signals as a function of the number of condensates in each *Nir*-47R cell. A reduced NirFP signal was shown in cells that contain more condensates (Figure 4c). These data suggested that gene expression can be possibly regulated by the CAG-repeat-mediated condensation of cellular mRNAs.

We next wanted to study if these CAG repeats can be used as trans-acting effectors to potentially control the cellular condensation of endogenous target RNAs. To test this, we first in vitro synthesized F30-2d×Broccoli-containing 47R strands (named as **47R-cO** and **47R-cY**) that were tagged with a complementary sequence that hybridize with either an OxyS or *lacY* target RNA. A 23-nt- and 26-nt-long non-structural region in OxyS and *lacY* was respectively designed as the targeting domain, which secondary structures were pre-evaluated via Mfold and NUPACK software.^28,29^ To image the proposed RNA–RNA interactions, a secondary fluorogenic RNA aptamer (Pepper)^30^ was tagged to the target OxyS and *lacY* RNAs, i.e., **OxyS-Pep** and ***lacY*-Pep**. After mixing OxyS-Pep with 47R-cO (Figure 5a) or *lacY*-Pep with 47R-cY (Figure 5b), the fluorescence signals from target RNAs (Pepper channel) were clearly accumulated and colocalized with the 47R condensates (Broccoli channel). As a negative control, without attaching the complementary sequence, 47R strands can still form obvious condensates but without recruiting OxyS-Pep or *lacY*-Pep (Figure 5a, b).

**Figure 5.**
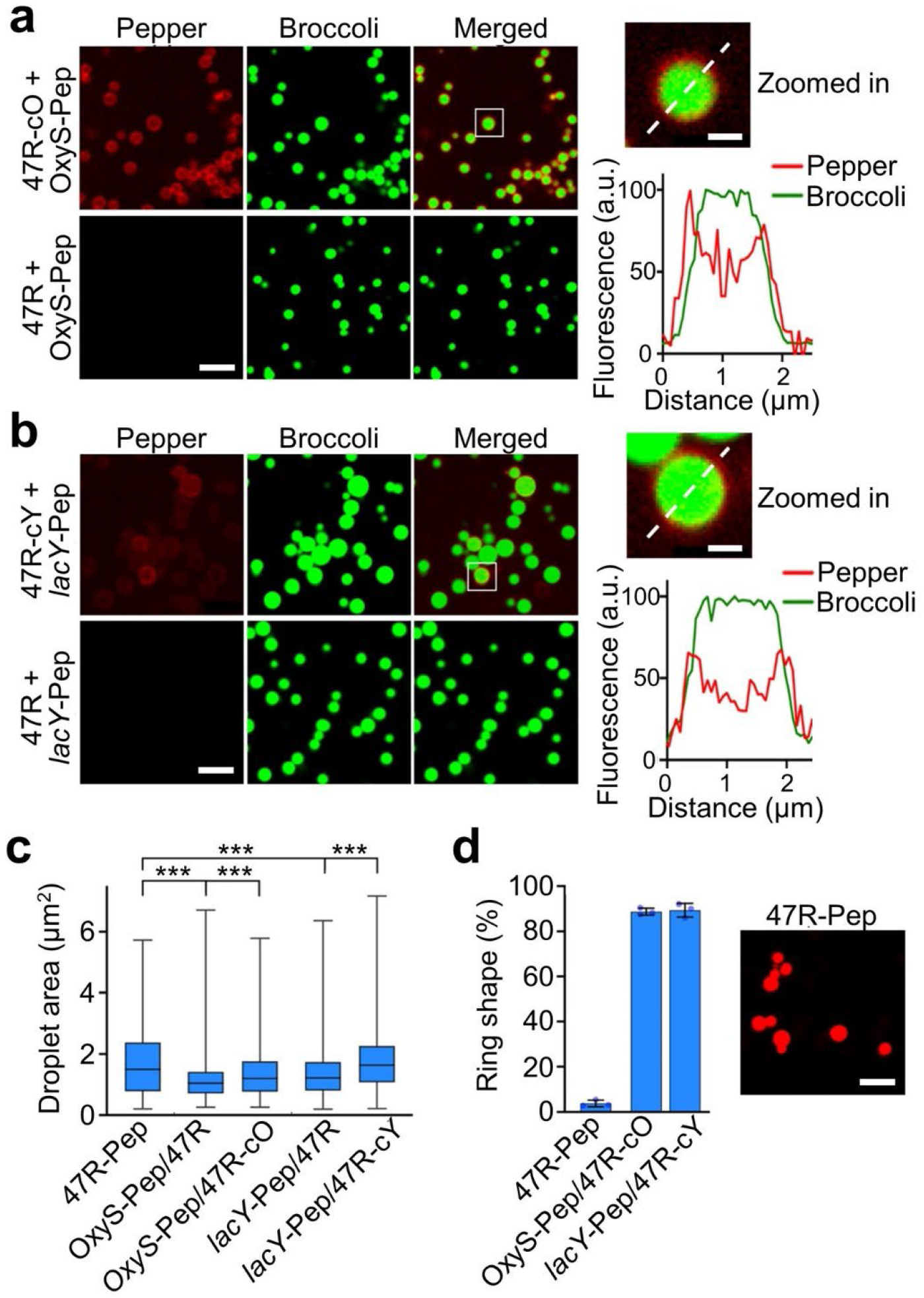
Targeted RNA recruitment into condensates via trans-acting CAG repeats. (**a, b**) Confocal fluorescence imaging of condensates induced by F30-2d×Broccoli-tagged 47×CAG repeats, with or without a complementary strand that target OxyS or *lacY* (47R-cO, 47R-cY, 47R). Pepper-tagged OxyS and *lacY* were used as target RNAs (OxyS-Pep, *lacY*-Pep). Insets show an enlarged view of condensate. Broccoli (green) and Pepper (red) fluorescence were plotted based on distance across the dashed line. Fluorescence images were taken in solutions containing 4 μM of each RNA, 20 mM MgCl_2_, 2 μM HBC620 and 80 μM DFHBI-1T after annealing. Scale bar, 5 μm. Inset scale bar, 1 μm. (**c**) Droplet area of each type of RNA condensate is plotted in box and min-to-max whiskers. (**d**) The percentage of ring-shaped condensates was measured in OxyS-Pep/47R-cO and *lacY*-Pep/47R-cY solutions, as compared to that of 47R-Pep. A total of 780–1010 condensates were analyzed in each case. Fluorescence imaging of condensation of Pepper-tagged 47×CAG repeats formed in solutions containing 4 μM RNA, 20 mM MgCl_2_, and 2 μM HBC620. Scale bar, 5 μm. Shown are the mean and SD values. Data was collected from at least three representative images. Two-tailed student’s t-test: ***, *p*<0.001.

Interestingly, a ring-shaped Pepper fluorescence pattern was observed in both OxyS-Pep/47R-cO and *lacY*-Pep/47R-cY condensate samples (Figure 5a, b). Meanwhile, the recruitment of target RNAs led to an increase in the condensate size (Figure 5c). These results suggested that the target RNAs were mainly hybridized to the surface areas of condensates, likely after the initial formation of the condensate core regions. To further assess if this ring-shaped fluorescence distribution could be resulted from the misfolding of Pepper RNAs inside the center of condensates, we synthesized a control strand with Pepper directly tagged with a 47×CAG-repeat (**47R-Pep**). 47R-Pep exhibited minimal ring-shaped condensates (Figure 5d), indicated that the ring-shaped Pepper fluorescence in OxyS-Pep/47R-cO and *lacY*-Pep/47R-cY samples were indeed likely due to the surface attachment of these target RNAs onto the CAG-repeat condensates.

We lastly tested if these trans-acting CAG-repeat tags can also function inside living cells. For this purpose, we transformed BL21 Star™ (DE3) cells with vectors expressing Pepper-tagged *NirFP* mRNA (***Nir-Pep***) together with an F30-2d×Broccoli-conjugated 47×CAG-repeat strand (47R) or that contains a 25-nt-long complementary sequence **47R-cN**. In the presence of 5 mM MgCl_2_, based on the Pearson’s correlation coefficient of the two fluorescent channels, ~80% of Pepper signals in *Nir*-Pep- and 47R-cN-expressing cells were colocalized with CAG-repeat condensates (Figure 6a, b). While in contrast, only 55% of *Nir*-Pep RNAs were found inside 47R condensates, indicating the roles of complementary sequence in recruiting target RNAs into CAG-repeat condensates. Consistent with the results from cis-acting CAG-repeat tags (Figure 4c), after forming condensates, ~30% lower NirFP signals were observed in 47R-cN/*Nir*-Pep cells as compared to that of 47R/*Nir*-Pep control (Figure 6c). All these data suggested that trans-acting CAG-repeat tags can be used to recruit cellular mRNAs into condensates, which may be potentially engineered into a functional gene regulation platform.

**Figure 6.**
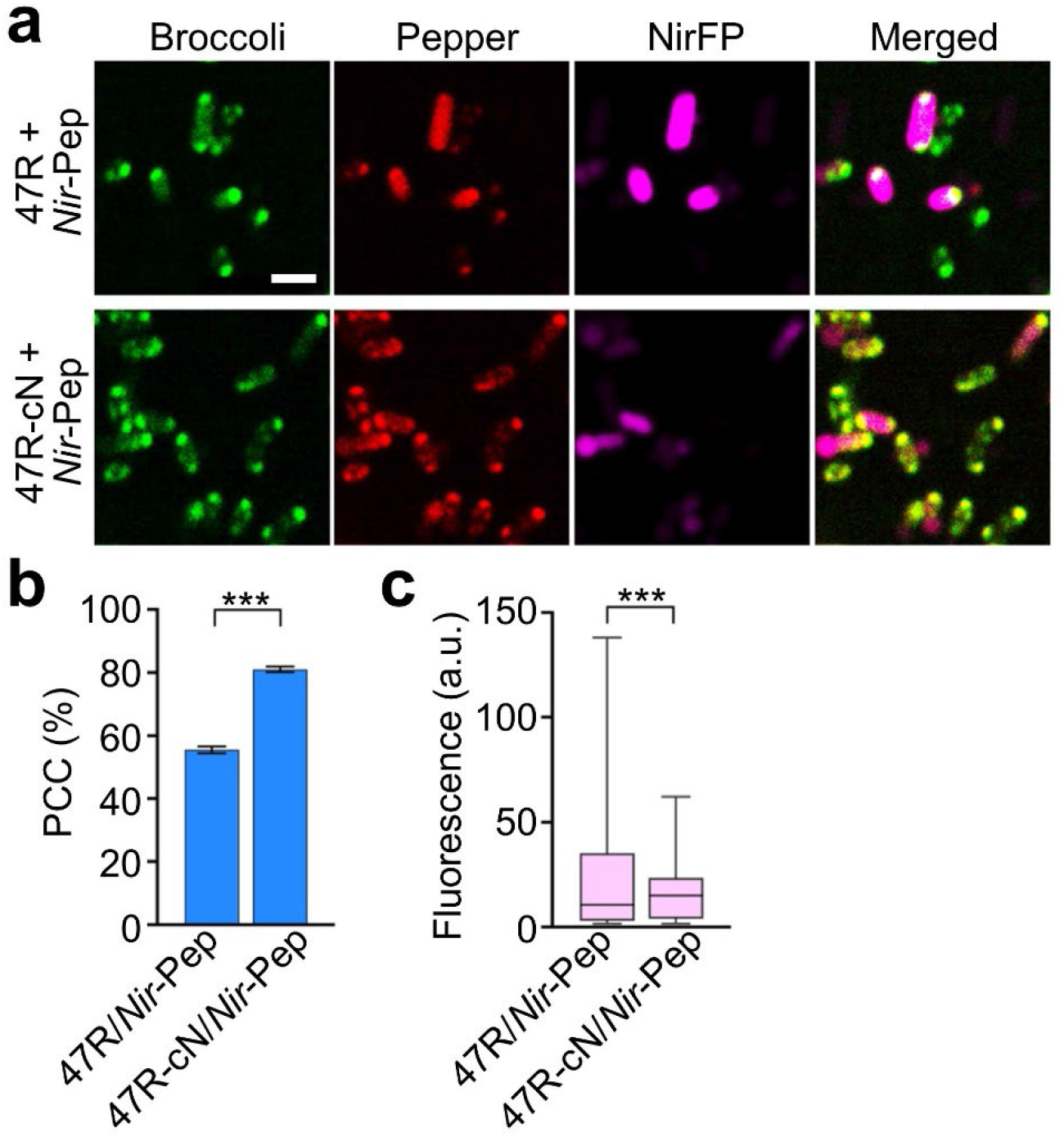
Trans-acting recruitment and functional regulation of target RNAs. (**a**) Confocal fluorescence imaging of *E. coli* cells that express Pepper-tagged near-infrared fluorescent protein RNA (*Nir*-Pep) and F30-2d×Broccoli-tagged 47×CAG repeats with or without a complementary targeting pair (47R-cN, 47R). Scale bar, 2 μm. (**b**) The Pearson’s correlation coefficient (PCC) of the Broccoli and Pepper fluorescence channels was quantified for both 47R and 47R-cN cells in the presence of 1 μM HBC620 and 5 mM Mg^2+^. A total of 1287 and 1764 cells were analyzed in each case. Shown are the mean and SEM values. (**c**) Average cellular NirFP fluorescence as measured in individual 47R or 47R-cN cells are plotted in box and min-to-max whiskers. A total of 717 and 742 cells were analyzed in each case. Data were collected from at least three representative images. Two-tailed student’s t-test: ***, *p*<0.001.

## Conclusions

The importance of RNA condensation in studying cellular functions as well as disease diagnosis and treatment has been increasingly recognized.^11,31^ Programmable and self-functional probes that enable precise cellular RNA condensate regulation are thus useful tools in the field of chemical biology and synthetic biology. In this study, we demonstrated that naturally existing CAG repeats can be used as genetically encoded tags to induce cellular condensation of different RNAs of interest, including small noncoding RNAs and long mRNAs (>3,000 nt). The formation of these self-assembled RNA condensates can be tracked *in vitro* and in living cells by fluorogenic RNA aptamers. The number, size, and density of RNA condensates can be facilely regulated by the length of CAG-repeat tags and Mg^2+^ concentration. The cellular RNA localization and compartmentalization can be rationally tuned by these CAG repeats tags, via either cis- or trans-acting mechanism. Critically, the cellular functions of these target RNAs such as in RNA–protein interactions (in the case of *lacY*) and gene expression patterns (in the case of *NirFP* and OxyS), can also be regulated by these CAG-repeat tags.

We expect that these functional CAG-repeat tags can be broadly applied to reprogram living cells with defined compartmentalizations and structures. This work can also inspire the potential engineering of various types of genetically encodable RNA nanodevices, which can be based on similar nature-inspired RNA repeat tags. These tags will be used to regulate different endogenous RNA targets and may also be applied to orthogonally recruit different RNA and protein molecules into specific RNA condensates. Powerful nanodevices may also be possibly designed to be controllable by different small-molecule or light triggers.

## Data Availability

The data found in this study are available within the main text and in the ESI online.

## Acknowledgements

The authors gratefully acknowledge the support from a UMass Amherst start-up grant, NSF CAREER award #1846152, Sloan Research Fellowship, and Camille Dreyfus Teacher-Scholar Award to M. You. R. Zheng was also supported by NIH T32GM008515. We are grateful to Dr. James Chambers for the assistance in fluorescence imaging. The authors also thank other members in the You Lab for useful discussion and valuable comments.

