## Supplementary Information for "Targeted RNA Condensation in Living Cells via Genetically Encodable Triplet Repeat Tags"

**Materials and Methods**

**Reagents and apparatus.** DNA oligonucleotides used in this work were synthesized and purified by Integrated DNA Technologies (Coralville, IA) and W. M. Keck Oligonucleotide Synthesis Facility (Yale University School of Medicine). The detailed sequences have been listed in Table S1. PCR products were cleaned using a Monarch® PCR & DNA Cleanup Kit [New England Biolabs (NEB), Ipswich, MA]. All the RNAs for in vitro experiments were transcribed using a HiScribe™ T7 high-yield RNA synthesis kit (New England Biolabs, Ipswich, MA) and purified with G-25 columns. All chemicals were of analytical grade and obtained from Sigma or Fisher Scientific unless otherwise noted. All the concentrations of nucleic acids were measured with a NanoDrop One UV-Vis spectrophotometer. The gel electrophoresis was performed on a BioRad electrophoresis analyzer (Bio-Rad, Hercules, CA) and imaged on a Bio-Rad Gel Doc EZ imager. Fluorescence measurements in solution were conducted with a BioTek Synergy 2 fluorescence plate reader ($\lambda_{ex}$= 485/20 nm, $\lambda_{em}$= 528/20 nm) (BioTek, Winooski, VT).

**Vector construction.** Different lengths of the CAG repeats were first cloned into a pAV-U6+27-F30-2xdBroccoli vector. In more detail, the vector was first digested with XbaI and SacII restriction enzymes (NEB). After purification via 1% agarose gel, the digested vector was ligated together with a similarly digested CAG repeat insert using T4 DNA ligase (NEB). The ligated product was then transformed into BL21 Star™ (DE3) cells (Thermo Fisher Scientific) and selected based on ampicillin resistance.

To construct CAG repeat-expressing pET-28c vectors for bacterial imaging, a pET-28c-F30-2xdBroccoli vector was digested with BglII and XhoI restriction enzymes (NEB), and then ligated together with a similarly digested CAG repeat insert using T4 DNA ligase. The CAG repeat inserts were synthesized by PCR using the above-prepared pAV-U6+27-F30-2xdBroccoli-CAG repeat vectors as the template and primers containing T7 terminator, and BglII and XhoI restriction sites. After T4 DNA ligation, the product was transformed into BL21 Star™ (DE3) cells and selected based on kanamycin resistance.

The OxyS, *lacY* and *lacZ* fragments were cloned respectively into pET-28c-F30-2xdBroccoli-CAG repeat vectors after digestion via BsaI and EcoRI restriction enzymes (NEB). The *lacY* and *lacZ* fragments were synthesized by PCR using the pYFP-lacY-1 and pSFV3-lacZ plasmids as the templates and primers that introduce BsaI and EcoRI restriction sites. After T4 DNA ligation, the product was transformed into BL21 Star™ (DE3) cells and selected based on kanamycin resistance. All these above-prepared plasmids were isolated using a GeneJET Plasmid Miniprep Kit (Thermo Fisher Scientific) and confirmed by Sanger sequencing (Eurofins Genomics).

***In vitro* RNA transcription and condensate formation.** For *in vitro* experiments, all the RNAs were transcribed using a HiScribe™ T7 high-yield RNA synthesis kit (NEB) and purified with G-25 columns. Template DNAs were prepared by PCR amplification from the above-mentioned pAV-U6+27- or pET-28c-based F30-2xdBroccoli-CAG repeat vectors. The *in vitro* formation of RNA condensates was prepared by first mixing 4 μM RNAs in buffer containing 10 mM Tris-HCl at pH=7.5, 100 mM KCl, and 20 mM $MgCl_2$. The mixture was then heated up to 95°C for 3 min and cooled down to 37°C at a rate of 2°C/min in a thermocycler. Afterwards, 80 μM DFHBI-1T was added and incubated for 15 min at 37°C before imaging using a Yokogawa spinning disk confocal on a Nikon Eclipse-TI inverted microscope. Images were collected with an excitation wavelength at 488 nm using a 100× oil immersion objective. The partition ratio of each RNA condensate was calculated based on the ratio of average fluorescence intensity within the condensate versus that in the solution region free of condensates. For the *in vitro* kinetic measurements, a solution containing 10 mM Tris-HCl at pH=7.5, 100 mM KCl, 20 mM $MgCl_2$, 4 μM RNA, and 80 μM DFHBI-1T was first heated up at 95°C for 3 min and then rapidly cooled down in ice for 30 sec right before starting to collect images.

**Cellular imaging.** RNA imaging in living bacterial cells was performed according to a previously established protocol.[1] Briefly, the BL21 Star™ (DE3) cells that express the corresponding RNAs were first grown in LB media at 37°C until the optical density at 600 nm ($OD_{600}$) reaches 0.4, and then 1 mM isopropyl-β-ᴅ-thiogalactoside (IPTG) was added for a 2-hour-induction. After the IPTG induction, the cells were adhered to poly-L-lysine-pretreated glass-bottom 8-well imaging plate (Cellvis, Mountain View, CA) in DPBS buffer for 45 min. Then the buffer was switched to fresh DPBS containing 200 μM DFHBI-1T and/or 1 μM HBC620 for a 30-min-incubation at 25°C before imaging. All the confocal fluorescence images were collected with a NiS-Elements AR software using a Yokogawa spinning disk confocal on a Nikon Eclipse-TI inverted microscope. Broccoli fluorescence signals were excited with a 488 nm laser, Pepper fluorescence signals were excited with a 561 nm laser, and the near-infrared fluorescent protein signal was collected under a 640 nm laser irradiation. A 100× oil immersion objective was used for collecting these cellular images. Structured illumination microscopy (SIM) super-resolution imaging was performed on a Nikon A1R-SIMe microscope equipped with a Hamamatsu sCMOS camera and a 100× oil immersion objective, under a 488 nm laser irradiation.

**Fluorescence recovery after photobleaching (FRAP).** The FRAP measurements were performed on an A1 spectral confocal microscope with a 100× oil immersion objective on a Nikon Eclipse-TI inverted system with an A1 stimulation module to ensure bleaching of a targeted area. Photobleaching was carried with a 488 nm laser for 1.0 s on a region of ~1 μm diameter. The fluorescence recovery after bleaching was then imaged every 5 s for a total of 3–5 min.

**Imaging data analysis.** Image analysis was performed using a NiS-Elements AR Analysis software. The set of actions performed on imaging channels were built as analysis recipes in the General Analysis 3 (*GA3*) module. For *in vitro* fluorescence images, a fluorescence intensity "threshold" $F_B$+3SD (background plus 3-fold of standard deviations on the 488 nm channel) and a diameter "threshold" of 0.5 μm was set to automatically detect each condensate. For each identified condensate, the "mean object intensity" and "object area" actions were applied to measure their corresponding mean fluorescence intensity and the area. The number of identified objects in one image (imaging view 4,430 $\mu m^2$) equals to the measured density of condensates. An inverted threshold was also set for measuring mean background intensities for the calculation of partition ratios.

For bacterial fluorescence images, after applying a "smooth" action, a fluorescence intensity "threshold" $F_B$+3SD (whole well background plus 3-fold of standard deviations on the 488 nm channel) was set to automatically detect each individual *E. coli* cells. The averaged whole cell fluorescence was measured via the "mean object intensity" action. To detect intracellular foci, the “bright spots” action was applied based on a fluorescence intensity “threshold” $F_B$+3SD (cellular background plus 3-fold of standard deviations on the 488 nm channel) and a diameter “threshold” of 0.5 μm. “Contrast” and “grow” were also set to enable the proper foci detection. Meanwhile, the “child ID” action was applied to set the detected cells as “parent” and the detected foci as “child” and chose “child is inside parent” condition. The count of foci in each detected cell was then obtained using the “group records” and “aggregate rows” actions. The “subtract” action was used to subtract the detected foci region from the detected cells region, i.e., the cellular background region. Then, the “mean object intensity” action was applied on both foci region and cellular background region to measure their corresponding mean fluorescence intensities and the partition ratio of each cellular condensate.

To further avoid inappropriate detection of cells and foci, only rod-shaped singly detected cells with properly detected foci were picked and also manually verified. Results shown in this study were generated by at least 90 above-verified cells from at least three representative images unless mentioned otherwise. At least two replicated experiments were performed for all these measurements. All the data analysis and fitting were performed using the ImageJ/FIJI and GraphPad Prism 9.2.0 software. Two-tailed student’s t-test was used to determine the statistical significance.

### Supplementary Figures

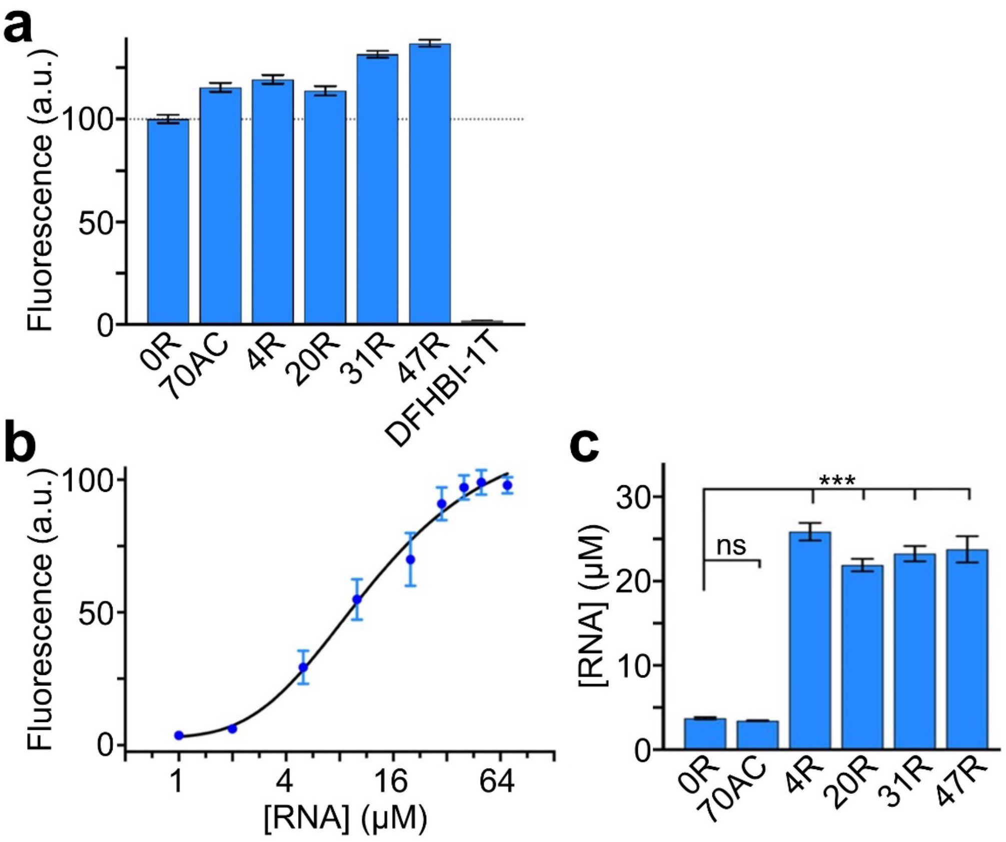


**Figure S1.** (**a**) Fluorescence measurement as performed in solutions containing 4 μM corresponding RNA, 20 mM $MgCl_2$, and 80 μM DFHBI-1T after annealing for the *in vitro* RNA condensate formation. F30-2d×Broccoli-tagged RNA strands containing 0×, 4×, 20×, 31×, 47×CAG repeats or 70×AC repeats are depicted as 0R, 4R, 20R, 31R, 47R, and 70AC, respectively. Shown are the mean and standard deviation (SD) values from three replicated experiments. (**b**) The correlation between RNA concentration and fluorescence signal intensities as measured in solutions containing 0–70 μM F30-2d×Broccoli RNA, 20 mM $MgCl_2$, and 80 μM DFHBI-1T under the same imaging condition for the *in vitro* RNA condensation characterization. Shown are the mean and SD values from three replicated experiments. (**c**) Based on the calibration curve shown in the panel (b), the estimated RNA concentration within each condensate formed in Figure 1. Shown are the mean and the standard error of the mean (SEM) values from three representative images. Two-tailed student's t-test: ***, $p<0.001$; ns, not significant, $p>0.05$.

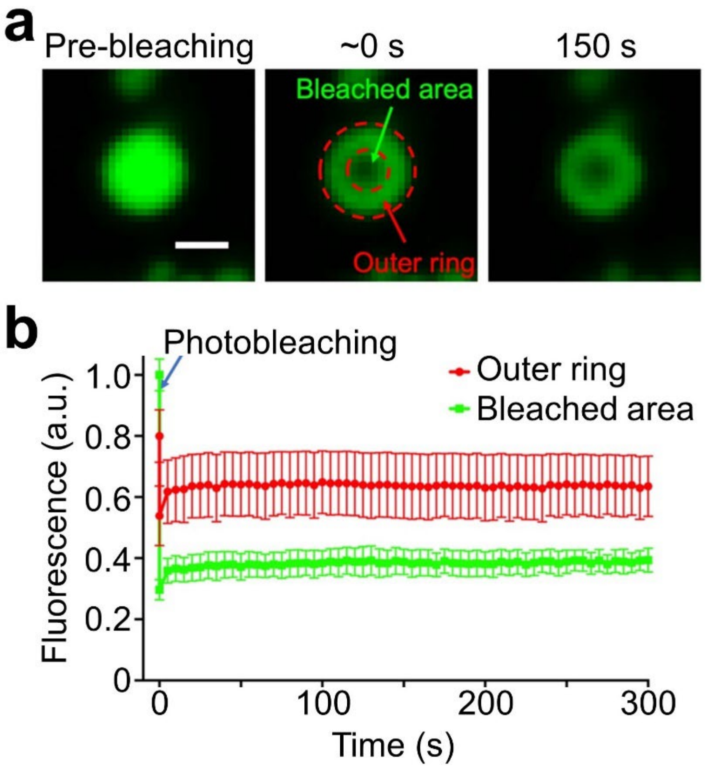


**Figure S2.** (**a**) *In vitro* fluorescence recovery after photobleaching (FRAP) measurement within condensates formed in solutions containing 4 μM 47R RNA, 20 mM $MgCl_2$, and 80 μM DFHBI-1T. Green and red arrow indicate the bleached area and outer ring region, respectively. Scale bar, 1 μm. (**b**) Averaged fluorescence intensity as measured within the bleached area (green line) and the outer ring region (red line) and plotted over time for the FRAP measurement. Shown are the mean and SEM values from at least three representative condensate FRAP measurements.

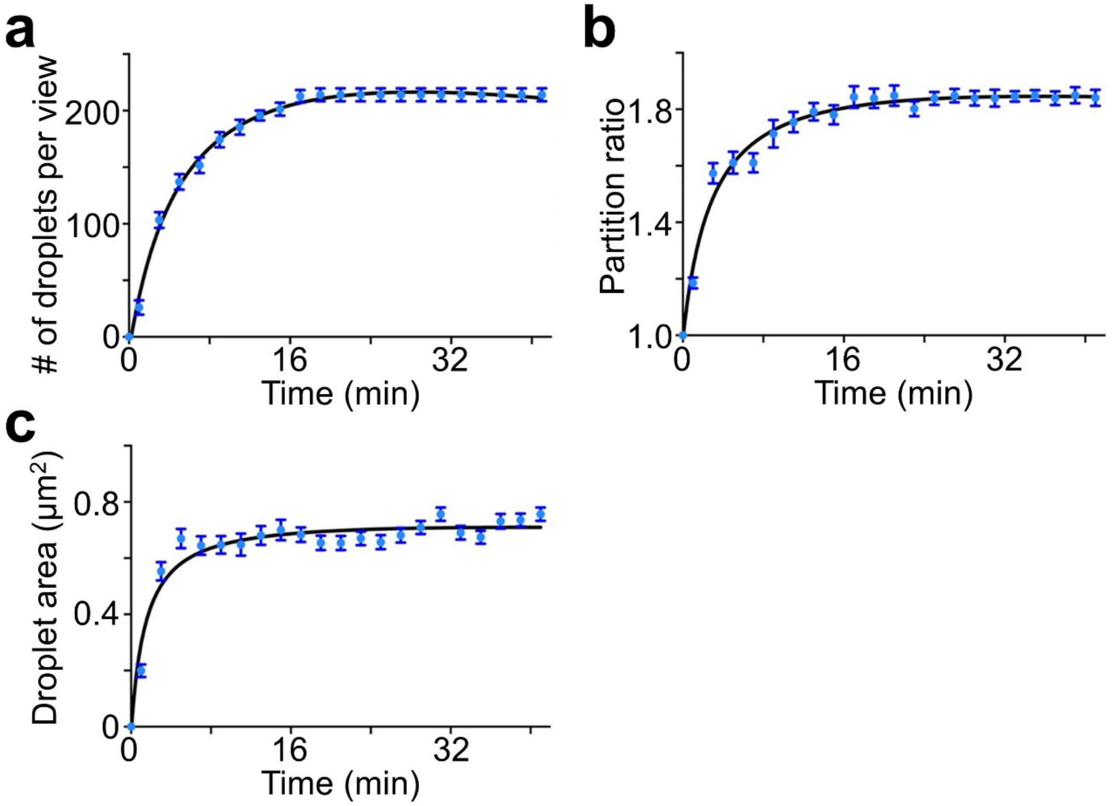


**Figure S3.** *In vitro* kinetic measurement of the condensate formation. (**a**) The number of condensates per imaging view, (**b**) the partition ratio and (**c**) area of each condensate are plotted after a 3 min 95°C heating and 30 s ice cooling. The condensate solution contains 4 µM 47R RNA, 20 mM $MgCl_2$, and 80 µM DFHBI-1T. The partition ratio is defined as the ratio of average fluorescence intensity inside individual condensate versus background signals in the solution. Shown are the mean and SEM values from at least three representative images, each imaging view equals to 4,430 µm$^2$.

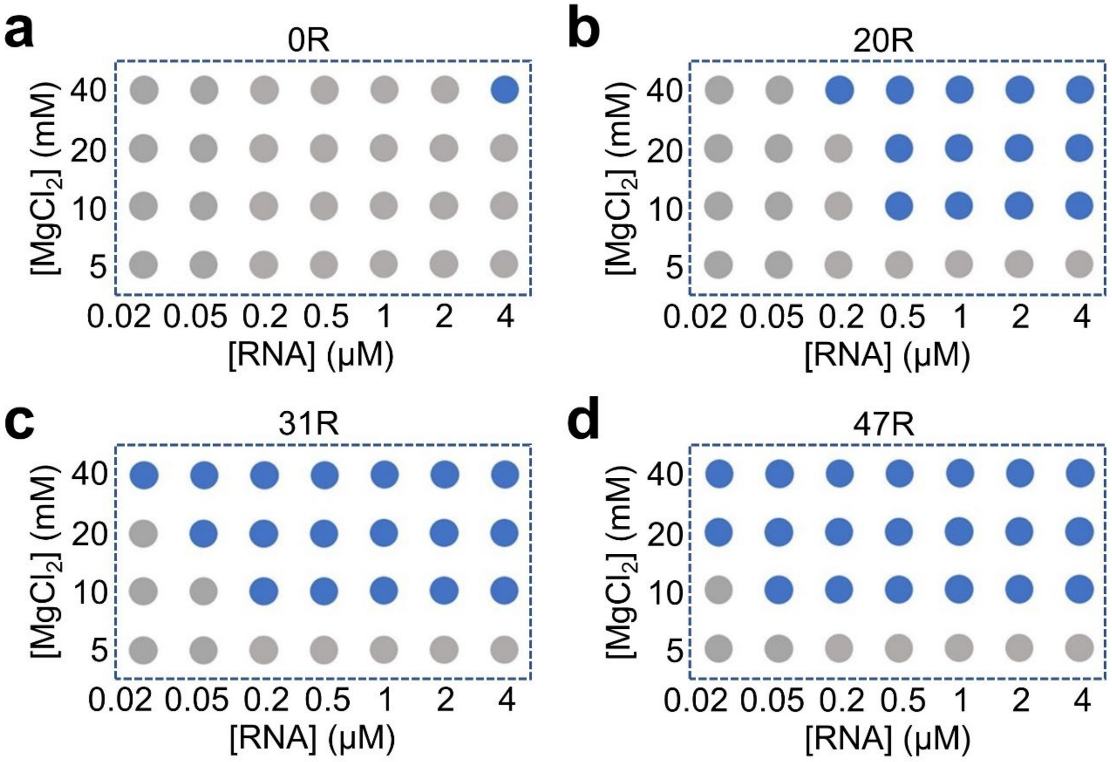


**Figure S4.** The phase diagrams of $Mg^{2+}$ and RNA concentration-dependent *in vitro* condensate formation. The measurements were performed in a solution after annealing for the condensate formation, which contains 10 mM Tris-HCl at pH=7.5, 100 mM KCl, and DFHBI-1T of 20-fold concentration as that of RNA. Blue (gray) dots represent the presence (absence) of >10 condensate with partition ratio >2 per 4,430 $\mu m^2$ imaging view from at least three replicated experiments.

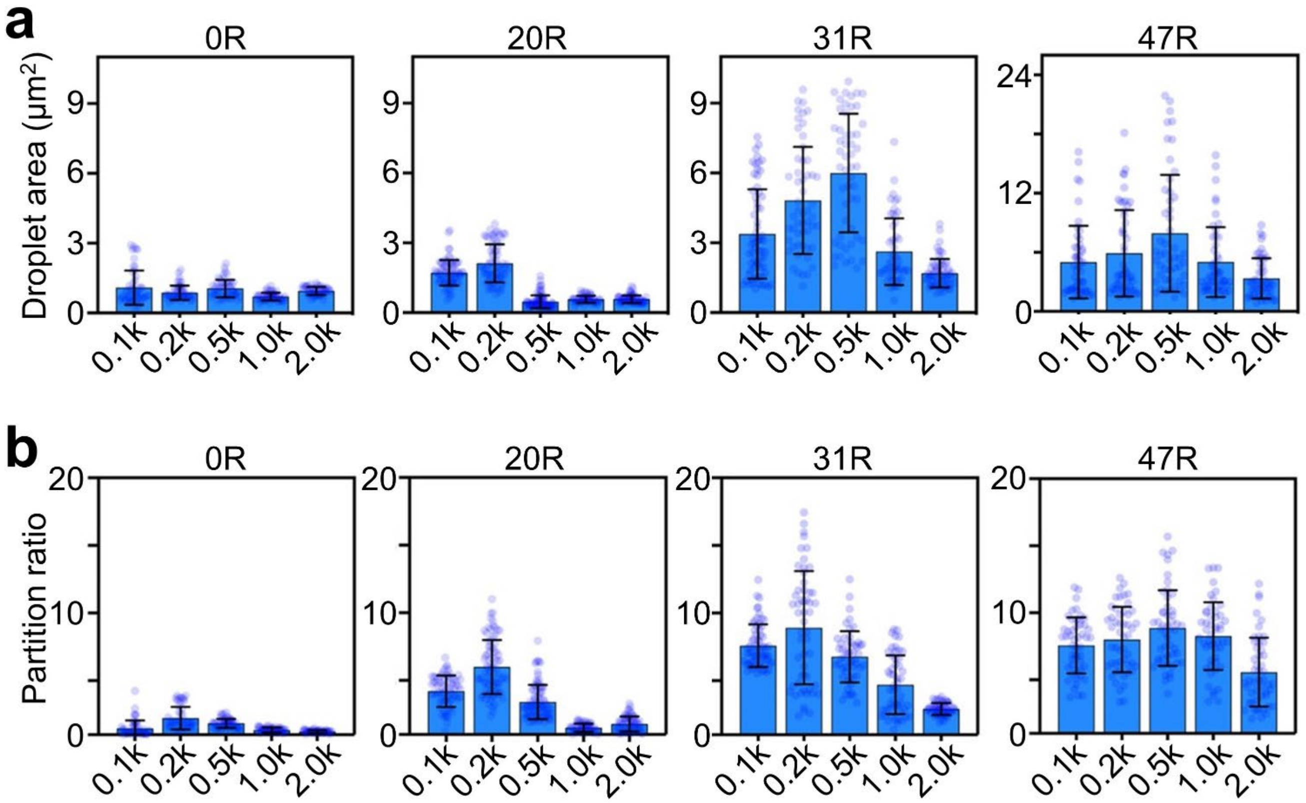


**Figure S5.** *In vitro* characterization of CAG-repeat-mediated condensation of different scrambled RNAs. (**a**) The area and (**b**) partition ratio of each condensate is plotted as a function of the length of scrambled target RNAs. The measurement was performed in solutions containing 4 μM corresponding RNA, 20 mM $MgCl_2$, and 80 μM DFH BI-1T after annealing for the condensate formation. Each data point represents one condensate. Shown are the mean and SD values from at least three representative images, each imaging view equals to 4,430 $\mu m^2$.

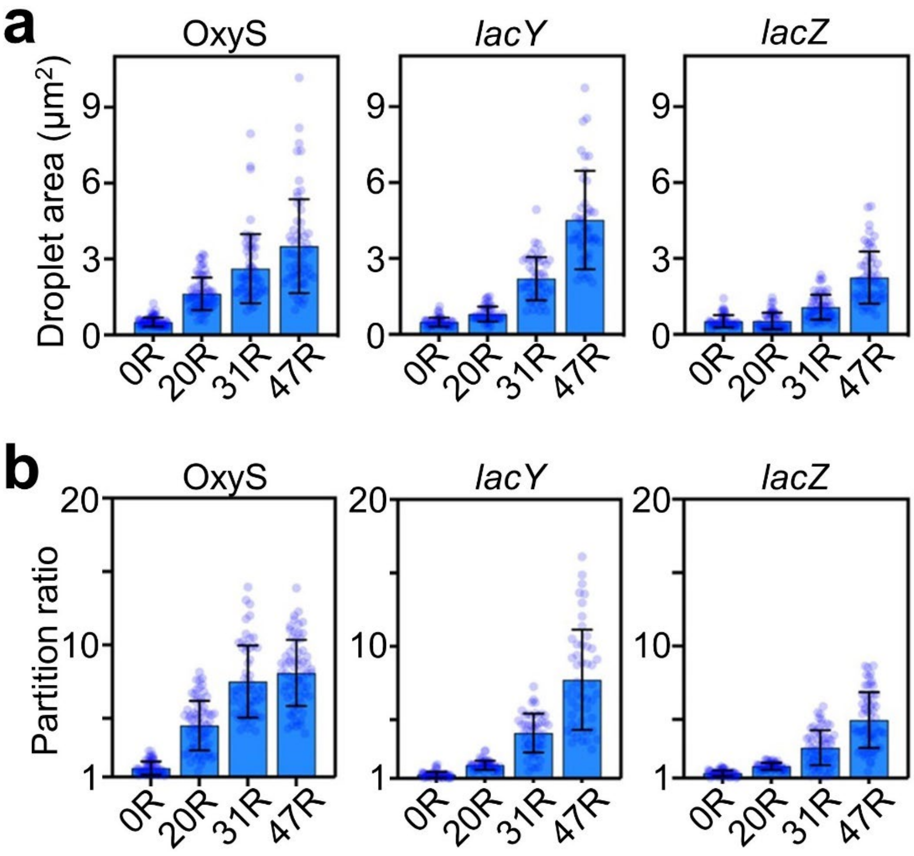


**Figure S6.** *In vitro* characterization of CAG-repeat-mediated condensation of different target RNAs. (**a**) The area and (**b**) partition ratio of each condensate is plotted as a function of the length of CAG repeats for each target RNAs. The measurement was performed in solutions containing 4 μM corresponding RNA, 20 mM $MgCl_2$, and 80 μM DFHBI-1T after annealing for the condensate formation. Each data point represents one condensate. Shown are the mean and SD values from at least three representative images, each imaging view equals to 4,430 $\mu m^2$.

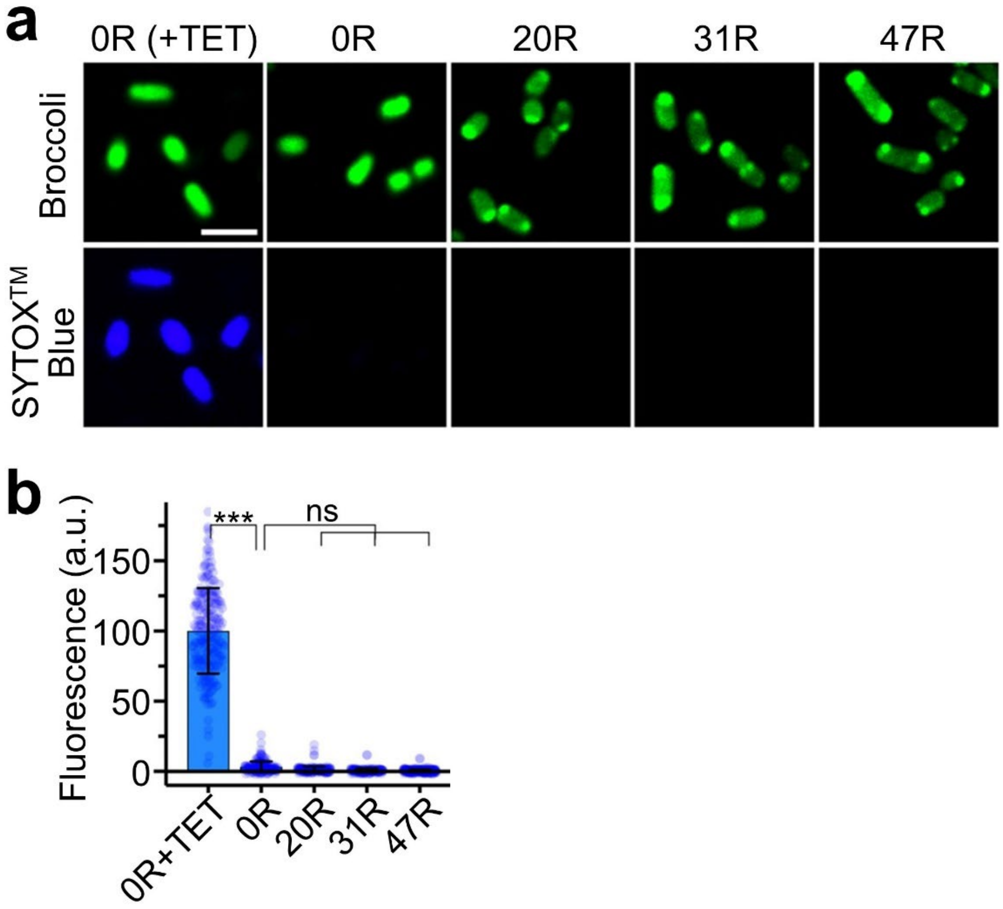


**Figure S7.** The cytotoxicity assessment of CAG-repeat-expressing bacterial cells. (**a**) Fluorescence imaging of BL21 Star™ (DE3) *E. coli* cells that express a pET-28c-F30-2d×Broccoli-0×, 20×, 31× or 47×CAG (0R, 20R, 31R or 47R) plasmid. These cells were incubated in DPBS buffer containing 1 μM SYTOX™ Blue for 2 h before imaging. As a positive control, one well of 0R cells were also treated with 1 mM tetracycline (TET) during the 2 h incubation with the SYTOX™ Blue dye. Scale bar, 3 μm. (**b**) Average cellular SYTOX™ Blue fluorescence as measured in individual cells. Each data point represents one cell. Shown are the mean and SD values. All the data is collected from at least three representative images. Two-tailed student's t-test: ***, $p<0.001$; ns, not significant, $p>0.05$.

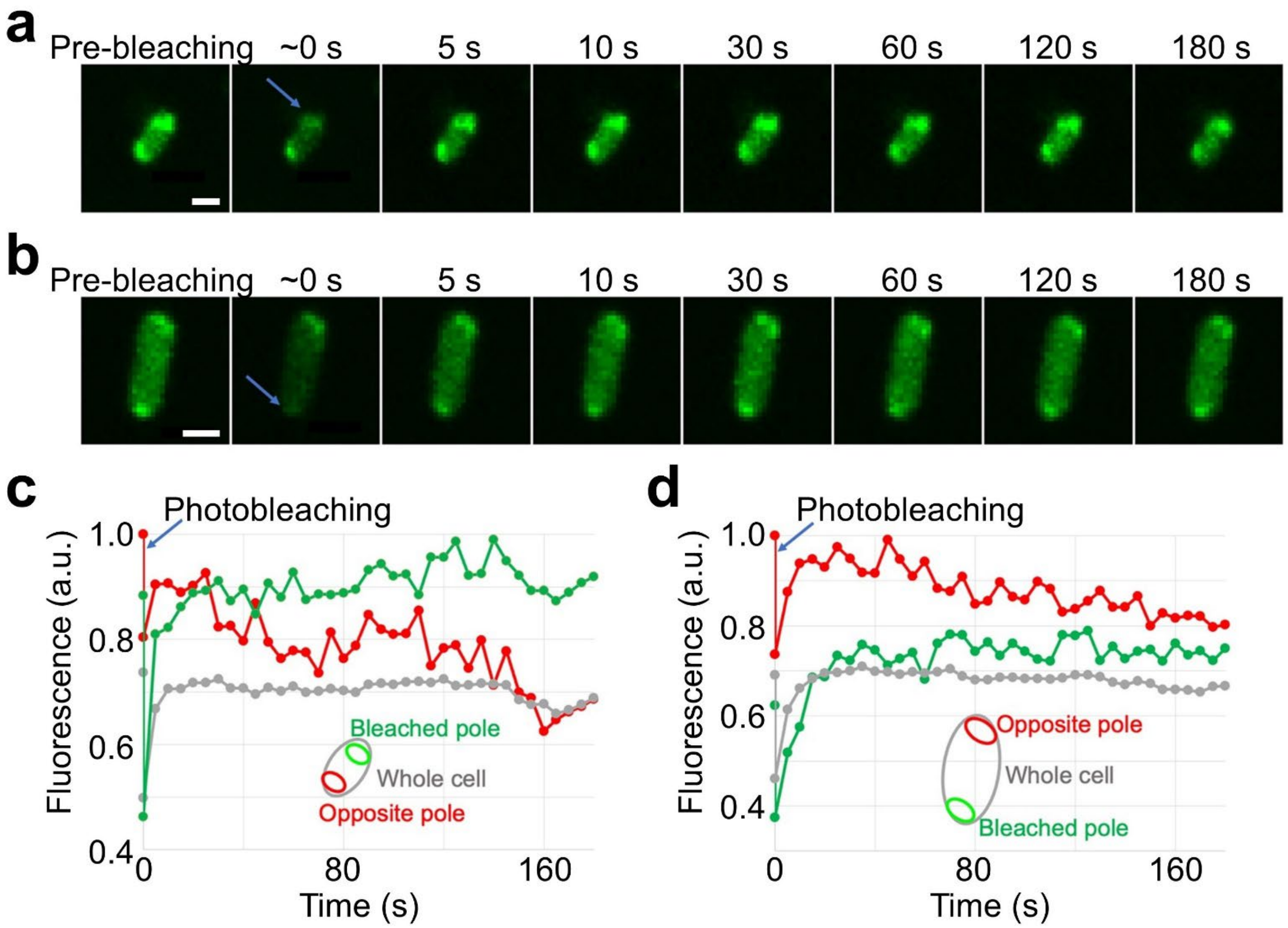


**Figure S8.** The FRAP measurement in BL21 Star$^{TM}$ (DE3) *E. coli* cells that express a pET-28c-F30-2d×Broccoli-47×CAG plasmid (47R). (**a**, **b**) Representative FRAP images from five out of nine measured cells containing two condensates at opposite poles. All these five cells show the transfer of fluorescence signals between two poles after photobleaching. Blue arrows indicate the bleached pole. Scale bar, 1 μm. (**c**, **d**) Averaged fluorescence intensity from the bleached pole (green dot/line), unbleached opposite pole (red dot/line) and whole cell (grey dot/line) are plotted over time for the FRAP measurement. The insets illustrate the corresponding measured regions.

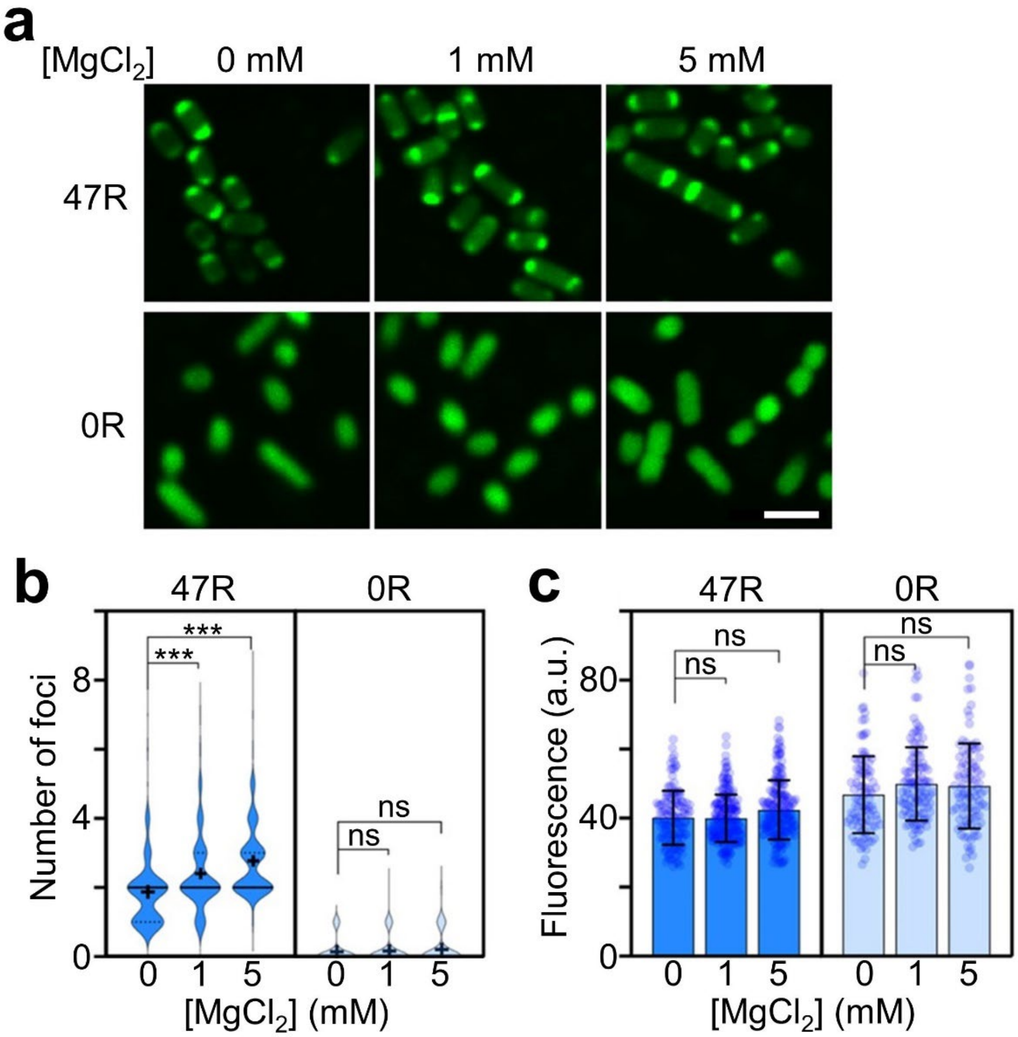


**Figure S9.** $Mg^{2+}$ concentration-regulated cellular RNA condensate formation. (**a**) Fluorescence imaging of BL21 Star™ (DE3) *E. coli* cells that express a pET-28c-F30-2d×Broccoli-0×CAG (0R) or pET-28c-F30-2d×Broccoli-47×CAG (47R) plasmid. These cells were incubated in DPBS buffer containing 0, 1, or 5 mM $MgCl_2$ for 2 h before imaging. Scale bar, 2 μm. (**b**) The violin plot distribution of the number of foci per cell, as measured from 0R or 47R cells. Solid and dashed line indicates the median and interquartile value, respectively. The black cross indicates the mean value. (**c**) The averaged fluorescence intensity as measured in each individual cell. Each data point represents one cell. Shown are the mean and SD values. All the data is collected from at least three representative images. Two-tailed student's t-test: ***, $p<0.001$; ns, not significant, $p>0.05$.

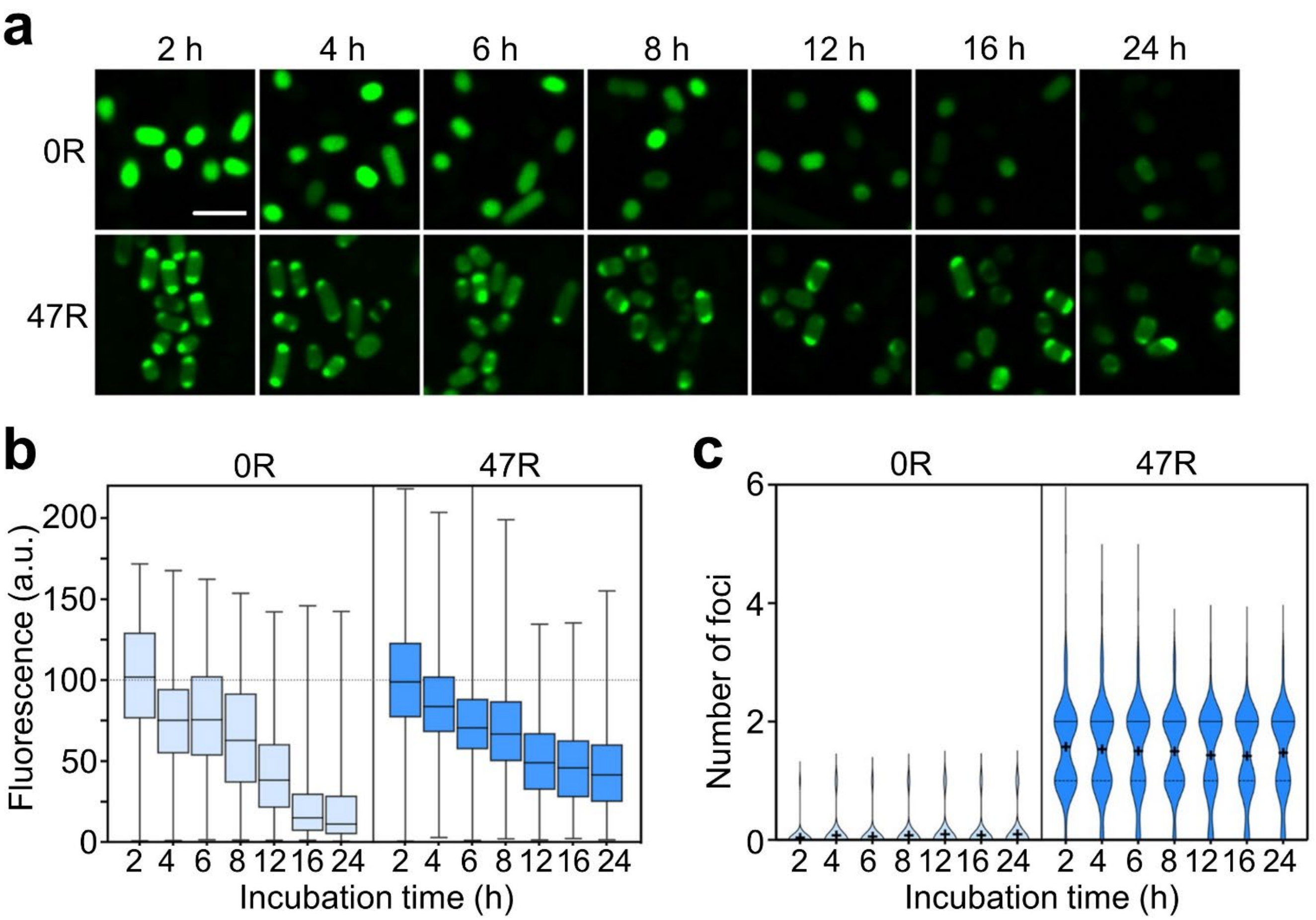


**Figure S10.** Condensate formation-mediated changes in cellular RNA lifetime. (**a**) Fluorescence imaging over 24 hours of BL21 Star™ (DE3) *E. coli* cells that express a pET-28c-F30-2d×Broccoli-0×CAG (0R) or pET-28c-F30-2d×Broccoli-47×CAG (47R) plasmid. These cells were first IPTG-induced for 2 h and then after removing the IPTG, left in the DPBS buffer for different time before imaging. Scale bar, 3 μm. (**b**) The averaged fluorescence intensity as measured in each individual cell. Shown are box plots with min-to-max whiskers collected from at least three representative images. (**c**) The violin plot distribution of the number of foci per cell, as measured from 0R or 47R cells after different time of incubation in the DPBS buffer. Solid and dashed line indicates the median and interquartile value, respectively. The black cross indicates the mean value. All the data is collected from at least three representative images.

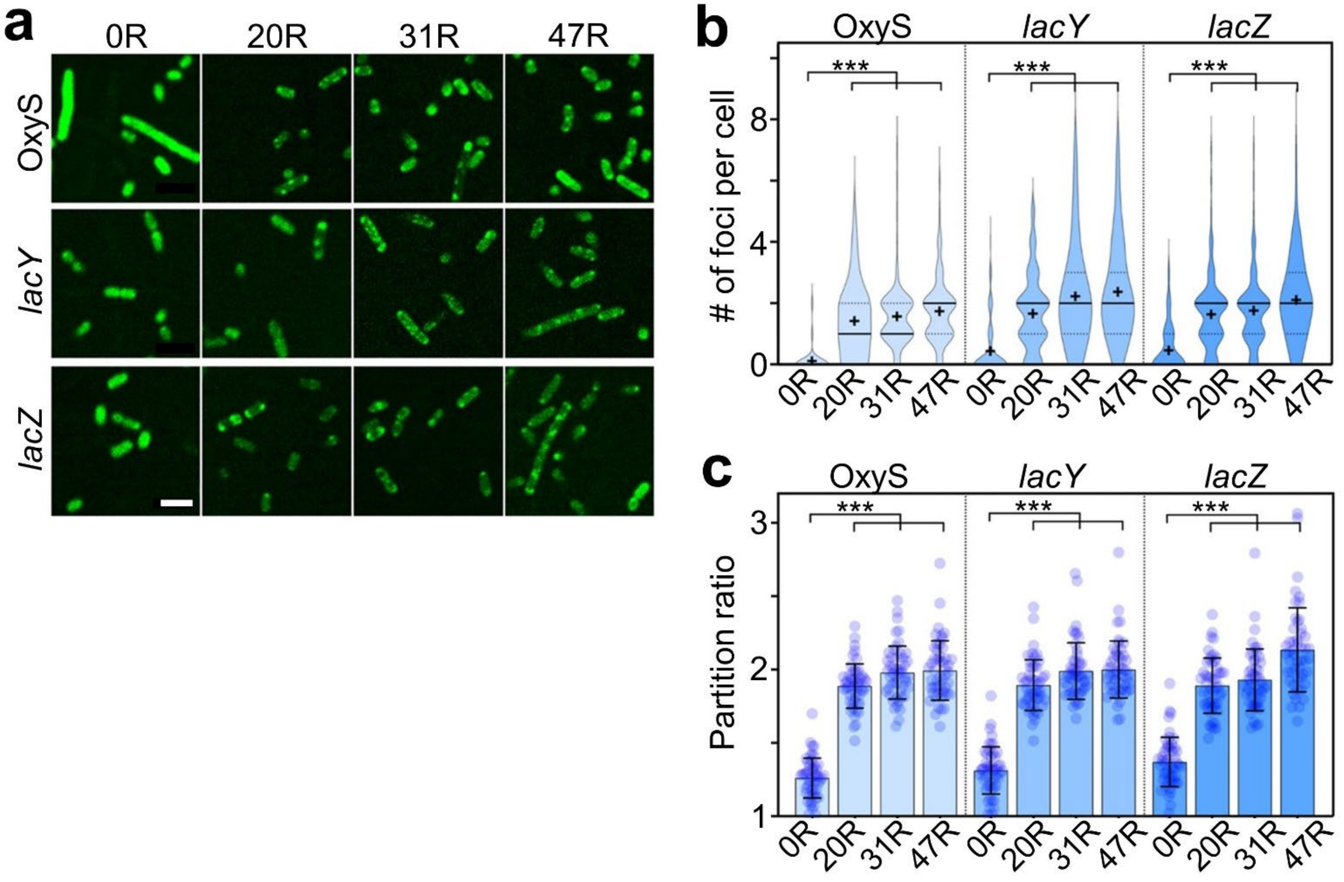


**Figure S11.** Condensation of target RNAs in *E. coli* cells. (**a**) Fluorescence imaging of BL21 Star™ (DE3) *E. coli* cells that express a pET-28c vector encoding 0R-, 20R-, 31R-, or 47R-tagged OxyS, *lacY* or *lacZ* target RNAs. Scale bar, 5 µm. (**b**) The violin plot distribution of the number of foci per cell, as measured from corresponding cells. Solid and dashed line indicates the median and interquartile value, respectively. The black cross indicates the mean value. (**c**) The partition ratio of individual cellular foci as measured from the same batch of cells. Each data point represents one cellular condensate. Shown are the mean and SD values. All the data is collected from at least three representative images. Two-tailed student's t-test with Bonferroni correction: ***, $p < 0.0003$.

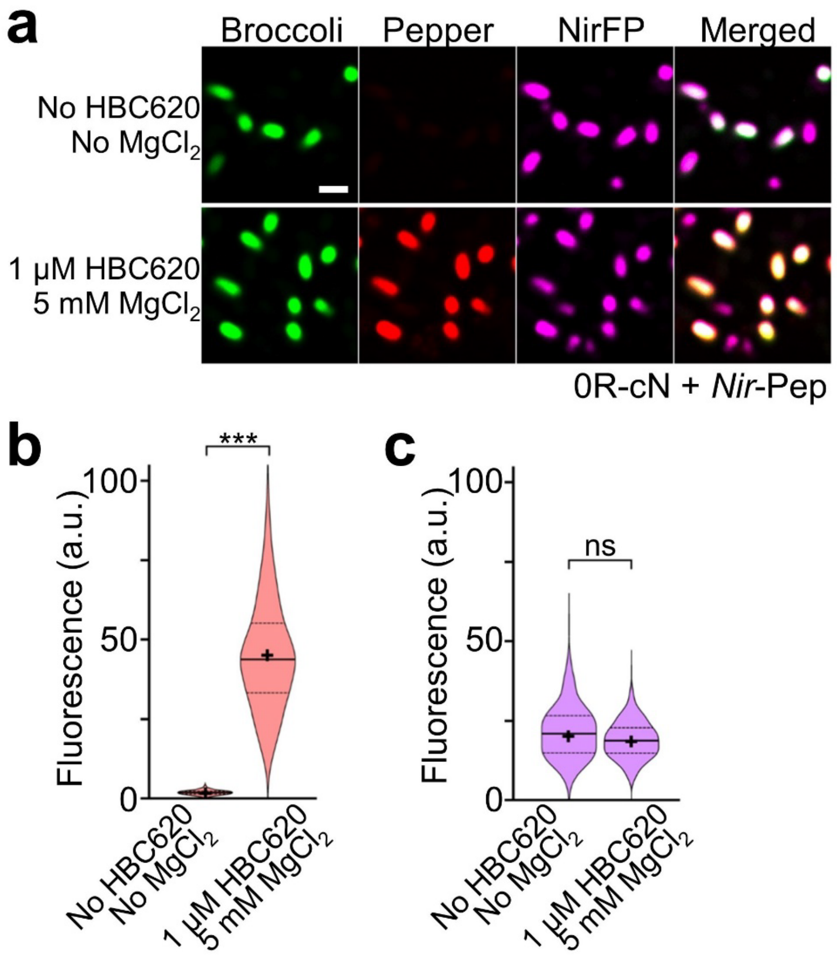


**Figure S12.** Trans-acting recruitment and regulation of target cellular RNAs. (**a**) Fluorescence imaging of BL21 Star™ (DE3) *E. coli* cells that express a pET-28c vector encoding Pepper-tagged near-infrared fluorescent protein (*Nir*-Pep) mRNA and a complementary strand-tagged 0R. Shown are the images before and after adding 1 μM HBC620 and 5 mM $Mg^{2+}$ in DPBS buffer. Scale bar, 2 μm. (**b**) Average cellular Pepper and (**c**) NirFP fluorescence signals as measured in 366 and 512 individual cells were plotted before and after adding 1 μM HBC620 and 5 mM $Mg^{2+}$, respectively. Shown are violin plots with solid and dashed line indicating the median and interquartile value, respectively. The black cross indicates the mean value. Data was collected from at least three representative images. Two-tailed student's t-test: ***, $p<0.001$; ns, not significant, $p>0.05$.

**Supplementary Table**

**Table S1.** Sequences of RNAs used in this study. Sequences for the F30-2d×Broccoli, Pepper and near-infrared fluorescent protein (*NirFP*) are shown in green, red and magenta, respectively. The CAG or AC repeating sequences are bolded and the complementary sequences for trans-acting are underlined.

| **RNA name** | **Sequences (5'-3')** |
|---|---|
| F30-2d×<br>Broccoli | UUGCCAUGUGUAUGUGGGAGACGGUCGGGUCCAUCUGAGACGGUCGGGUCCAGAUAUUCGUAUC<br>UGUCGAGUAGAGUGUGGGCUCAGAUGUCGAGUAGAGUGUGGGCUCCCACAUACUCUGAUGAUCC<br>AGACGGUCGGGUCCAUCUGAGACGGUCGGGUCCAGAUAUUCGUAUCUGUCGAGUAGAGUGUGGG<br>CUCAGAUGUCGAGUAGAGUGUGGGCUGGAUCAUUCAUGGCAA |
| 4R | F30-2d×Broccoli-UCUAGAGAAUUCGGUCUCA**CAGCAGCAGCAG**UUUUUUCCGCGG |
| 20R | F30-2d×Broccoli-UCUAGAGAAUUCGGUCUCA**CAGCAGCAGCAGCAGCAGCAGCAGCAGCAGC<br>AGCAGCAGCAGCAGCAGCAGCAGCAG**UUUUUUCCGCGG |
| 31R | F30-2d×Broccoli-UCUAGAGAAUUCGGUCUCA**CAGCAGCAGCAGCAGCAGCAGCAGCAGCAGC<br>AGCAGCAGCAGCAGCAGCAGCAGCAGCAGCAGCAGCAGCAGCAGCAGCAGCAG**UUUU<br>UUUCCGCGG |
| 47R | F30-2d×Broccoli-UCUAGAGAAUUCGGUCUCA**CAGCAGCAGCAGCAGCAGCAGCAGCAGCAGC<br>AGCAGCAGCAGCAGCAGCAGCAGCAGCAGCAGCAGCAGCAGCAGCAGCAGCAGCAGC<br>AGCAGCAGCAGCAGCAGCAGCAGCAGCAGCAGCAGCAGCAG**UUUUUUCCGCGG |
| 70AC | F30-2d×Broccoli-UCUAGAGAAUUCGGUCUCA**ACACACACACACACACACACACACACACACACA<br>CACACACACACACACACACACACACACACACACACACACACACACACACACACACACACAC<br>ACACACACACACACACACACACACACACACACACACACACACA**UUUUUUCCGCGG |
| Pepper | GGAUCCCCAAUCGUGGCGUGUCGGCCUGCUUCGGCAGGCACUGGCGCCGGGAUCC |
| Scrambled<br>0.1k RNA | CCUUUACCUGUAGGUGCGAUACCAUCGUAUCAUGUGAAGGGUACGUAGUUAAGAAAUCACUAUG<br>UGCCCCGGCCUGUACGGUAAAACGGUAGGGUACGC |
| Scrambled<br>0.2k RNA | CUAGCAGGCGCGUAUCCAACCUACGCCACAAACUGGGCCGACGAGCAGGUGUUACAGGCCAGGAA<br>CAUAGGACUGUGUGCAGCAUCCUUGACUGAGGGAAGACUCGGCAAACUGUCCAUUCUCCGCAAGA<br>AGCAAUUGAAACCUUGCGACACAGUCAUGUUCUCGGUAGGAUCUACAUUGUACACUGAGAGCAGA<br>AAGCU |
| Scrambled<br>0.5k RNA | GAUGGCGGAUGUGUGACAUACACGACGCCAAAAGAUUUUGUUCCAGCUCCUGCCACCUCCGCUAC<br>GCGAGAGAUUAACCACCCACGAUGGCCGCCAAAGUGCAUGUUGAUAUUGAGGCUGACAGCCCAUU<br>CAUCAAGUCUUUGCAGAAGGCAUUUCCGUCGUUCGAGGUGGAGUCAUUGCAGGUCACACCAAAUG<br>ACCAUGCAAAUGCCAGAGCAUUUUCGCACCUGGCUACCAAAUUGAUCGAGCAGGAGACUGACAAA<br>GACACACUCAUCUUGGAUAUCGGCAGUGCGCCUUCCAGGAGAAUGAUGUCUACGCACAAAUACCA<br>CUGCGUAUGCCCUAUGCGCAGCGCAGAAGACCCCGAAAGGCUCGAUAGCUACGCAAAGAAACUGG<br>CAGCGGCCUCCGGGAAGGUGCUGGAUAGAGAGAUCGCAGGAAAAAUCACCGACCUGCAGACCGU<br>CAUGGCUACGCCAGACGCUGAAUCUCCUACCUUUUGCCUGCAUACA |
| Scrambled<br>1.0k RNA | CGUGACGUAUCACGCGGAGGGAUUCCUAGUGUGCAAGACCACAGACACUGUCAAAGGAGAAAGAG<br>UCUCAUUCCCUGUAUGCACCUACGUCCCCUCAACCAUCUGUGAUCAAAUGACUGGCAUACUAGCG<br>ACCGACGUCACACCGGAGGACGCACAGAAGUUGUUAGUGGGAUUGAAUCAGAGGAUAGUUGUGAA<br>CGGAAGAACACAGCGAAACACUAACACGAUGAAGAACUAUCUGCUUCCGAUUGUGGCCGUCGCAU<br>UUAGCAAGUGGGCGAGGGAAUACAAGGCAGACCUUGAUGAUGAAAAACCUCUGGGUGUCCGAGAG<br>AGGUCACUUACUUGCUGCUGCUUGUGGGCAUUUAAAACGAGGAAGAUGCACACCAUGUACAAGAA<br>ACCAGACACCCAGACAAUAGUGAAGGUGCCUUCAGAGUUUAACUCGUUCGUCAUCCCGAGCCUAU<br>GGUCUACAGGCCUCGCAAUCCCAGUCAGAUCACGCAUUAAGAUGCUUUUGGCCAAGAAGACCAAG<br>CGAGAGUUAAUACCUGUUCUCGACGCGUCGUCAGCCAGGGAUGCUGAACAAGAGGAGAAGGAGA<br>GGUUGGAGGCCGAGCUGACUAGAGAAGCCUUACCACCCCUCGUCCCCAUCGCGCCGGCGGAGAC |

| | |
|---|---|
| | GGGAGUCGUCGACGUCGACGUUGAAGAACUAGAGUAUCACGCAGGUGCAGGGGUCGUGGAAACA<br>CCUCGCAGCGCGUUGAAAGUCACCGCACAGCCGAACGACGUACUACUAGGAAAUUACGUAGUUCU<br>GUCCCCGCAGACCGUGCUCAAGAGCUCCAAGUUGGCCCCCGUGCACCCUCUAGCAGAGCAGGUG<br>AAAAUAAUAACACAUAACGGGAGGGCCGGCGGUUACCAGGUCGACGGAUAUGACGGCAGGGUCCU<br>ACUACCAUGUGGAUCGGCCAUUCCGGUCCCUGAGUUUCAAGCUUUGAGCGAGAGCGCCACUAUG<br>GUGUACAACGAAAGGGAGUUCGUCAACAGG |
| Scrambled 2.0k RNA | AAACUAUACCAUAUUGCCGUUCACGGACCGUCGCUGAACACCGACGAGGAGAACUACGAGAAAGU<br>CAGAGCUGAAAGAACUGACGCCGAGUACGUGUUCGACGUAGAUAAAAAAUGCUGCGUCAAGAGAG<br>AGGAAGCGUCGGGUUUGGUGUUGGUGGGAGAGCUAACCAACCCCCCGUUCCAUGAAUUCGCCUA<br>CGAAGGGCUGAAGAUCAGGCCGUCGGCACCAUAUAAGACUACAGUAGUAGGAGUCUUUGGGGUU<br>CCGGGAUCAGGCAAGUCUGCUAUUAUUAAGAGCCUCGUGACCAAACACGAUCUGGUCACCAGCGG<br>CAAGAAGGAGAACUGCCAGGAAAUAGUUAACGACGUGAAGAAGCACCGCGGGAAGGGGACAAGUA<br>GGGAAAACAGUGACUCCAUCCUGCUAAACGGGUGUCGUCGUGCCGUGGACAUCCUAUAUGUGGA<br>CGAGGCUUUCGCUUGCCAUUCCGGUACUCUGCUGGCCCUAAUUGCUCUUGUUAAACCUCGGAGC<br>AAAGUGGUGUUAUGCGGAGACCCCAAGCAAUGCGGAUUCUUCAAUAUGAUGCAGCUUAAGGUGAA<br>CUUCAACCACAACAUCUGCACUGAAGUAUGUCAUAAAAGUAUAUCCAGACGUUGCACGCGUCCAG<br>UCACGGCCAUCGUGUCUACGUUGCACUACGGAGGCAAGAUGCGCACGACCAACCCGUGCAACAAA<br>CCCAUAAUCAUAGACACCACAGGACAGACCAAGCCCAAGCCAGGAGACAUCGUGUUAACAUGCUUC<br>CGAGGCUGGGCAAAGCAGCUGCAGUUGGACUACCGUGGACACGAAGUCAUGACAGCAGCAGCAU<br>CUCAGGGCCUCACCCGCAAAGGGGUAUACGCCGUAAGGCAGAAGGUGAAUGAAAAUCCCUUGUAU<br>GCCCCUGCGUCGGAGCACGUGAAUGUACUGCUGACGCGCACUGAGGAUAGGCUGGUGUGGAAAA<br>CGCUGGCCGGCGAUCCCUGGAUUAAGGUCCUAUCAAACAUUCCACAGGGUAACUUUACGGCCACA<br>UUGGAAGAAUGGCAAGAAGAACACGACAAAAUAAUGAAGGUGAUUGAAGGACCGGCUGCGCCUGU<br>GGACGCGUUCCAGAACAAAGCGAACGUGUGUUGGGCGAAAAGCCUGGUGCCUGUCCUGGACACU<br>GCCGGAAUCAGAUUGACAGCAGAGGAGUGGAGCACCAUAAUUACAGCAUUUAAGGAGGACAGAGC<br>UUACUCUCCAGUGGUGGCCUUGAAUGAAAUUUGCACCAAGUACUAUGGAGUUGACCUGGACAGUG<br>GCCUGUUUUCUGCCCCGAAGGUGUCCCUGUAUUACGAGAACAACCACUGGGAUAACAGACCUGGU<br>GGAAGGAUGUAUGGAUUCAAUGCCGCAACAGCUGCCAGGCUGGAAGCUAGACAUACCUUCCUGAA<br>GGGGCAGUGGCAUACGGGCAAGCAGGCAGUUAUCGCAGAAAGAAAAAUCCAACCGCUUUCUGUGC<br>UGGACAAUGUAAUUCCUAUCAACCGCAGGCUGCCGCACGCCCUGGUGGCUGAGUACAAGACGGU<br>UAAAGGCAGUAGGGUUGAGUGGCUGGUCAAUAAAGUAAGAGGGUACCACGUCCUGCUGGUGAGU<br>GAGUACAACCUGGCUUUGCCUCGACGCAGGGUCACUUGGUUGUCACCGCUGAAUGUCACAGGCG<br>CCGAUAGGUGCUACGACCUAAGUUUAGGACUGCCGGCUGACGCCGGCAGGUUCGACUUGGUCUU<br>UGUGAACAUUCACACGGAAUUCAGAAUCCACCACUACCAGCAGUGUGUCGACCACGCCAUGAAGC<br>UGCAGAUGCUUGGGGGAGAUGCGCUACGACUGCUAAAACCCGGCGGCAUCUUGAUGAGAGCUUA<br>CGGAUACGCCGAUAAAAUCAGCGAAGCCGUUGUUUCCUCCUUAAGCAGAAAGUUCUCGUCUGCAA<br>GAGUGUUGCGCCCGGAUUGUGUCACCAGCAAUACAGAAGUGUUCUUGCUGUUCUCCAACUU |
| OxyS<br>(109 nt) | GAAACGGAGCGGCACCUCUUUUAACCCUUGAAGUCACUGCCCGUUUCGAGAGUUUCUCAACUCGA<br>AUAACUAAAGCCAACGUGAACUUUUGCGGAUCUCCAGGAUCCGC |
| *lacY*<br>(1254 nt) | AUGUACUAUUUAAAAAACACAAACUUUUGGAUGUUCGGUUUAUUCUUUUUCUUUUACUUUUUUAUC<br>AUGGGAGCCUACUUCCCGUUUUUCCCGAUUUGGCUACAUGACAUCAACCAUAUCAGCAAAAGUGA<br>UACGGGUAUUAUUUUUGCCGCUAUUUCUCUGUUCUCGCUAUUAUUCCAACCGCUGUUUGGUCUG<br>CUUUCUGACAAACUCGGGCUGCGCAAAUACCUGCUGUGGAUUAUUACCGGCAUGUUAGUGAUGUU<br>UGCGCCGUUCUUUAUUUUUAUCUUCGGGCCACUGUUACAAUACAACAUUUUAGUAGGAUCGAUUG<br>UUGGUGGUAUUUAUCUAGGCUUUUGUUUUAACGCCGGUGCGCCAGCAGUAGAGGCAUUUAUUGA<br>GAAAGUCAGCCGUCGCAGUAAUUUCGAAUUUGGUCGCGCGCGGAUGUUUGGCUGUGUUGGCUGG<br>GCGCUGUGUGCCUCGAUUGUCGGCAUCAUGUUCACCAUCAAUAAUCAGUUUGUUUUCUGGCUGG<br>GCUCUGGCUGUGCACUCAUCCUCGCCGUUUUACUCUUUUUCGCCAAAACGGAUGCGCCCUCUUC<br>UGCCACGGUUGCCAAUGCGGUAGGUGCCAACCAUUCGGCAUUUAGCCUUAAGCUGGCACUGGAA<br>CUGUUCAGACAGCCAAAACUGUGGUUUUUGUCACUGUAUGUUAUUGGCGUUUCCUGCACCUACGA<br>UGUUUUUGACCAACAGUUUGCUAAUUUCUUUACUUCGUUCUUUGCUACCGGUGAACAGGGUACGC<br>GGGUAUUUGGCUACGUAACGACAAUGGGCGAAUUACUUAACGCCUCGAUUAUGUUCUUUGCGCCA<br>CUGAUCAUUAAUCGCAUCGGUGGGAAAAACGCCCUGCUGCUGGCUGGCACUAUUAUGUCUGUAC |

| | |
|---|---|
| | GUAUUAUUGGCUCAUCGUUCGCCACCUCAGCGCUGGAAGUGGUUAUUCUGAAAACGCUGCAUAUG<br>UUUGAAGUACCGUUCCUGCUGGUGGGCUGCUUUAAAUAUAUUACCAGCCAGUUUGAAGUGCGUU<br>UUUCAGCGACGAUUUAUCUGGUCUGUUUCUGCUUCUUUAAGCAACUGGCGAUGAUUUUUAUGUCU<br>GUACUGGCGGGCAAUAUGUAUGAAAGCAUCGGUUUCCAGGGCGCUUAUCUGGUGCUGGGUCUGG<br>UGGCGCUGGGCUUCACCUUAAUUUCCGUGUUCACGCUUAGCGGCCCCGGCCCGCUUUCCCUGCU<br>GCGUCGUCAGGUGAAUGAAGUCGCUUAA |
| *lacZ*<br>(3048 nt) | GUCGUUUUACAACGUCGUGACUGGGAAAACCCUGGCGUUACCCAACUUAAUCGCCUUGCAGCACA<br>UCCCCCUUUCGCCAGCUGGCGUAAUAGCGAAGAGGCCCGCACCGAUCGCCCUUCCCAACAGUUG<br>CGCAGCCUGAAUGGCGAAUGGCGCUUUGCCUGGUUUCCGGCACCAGAAGCGGUGCCGGAAAGCU<br>GGCUGGAGUGCGAUCUUCCUGAGGCCGAUACUGUCGUCGUCCCCUCAAACUGGCAGAUGCACGG<br>UUACGAUGCGCCCAUCUACACCAACGUAACCUAUCCCAUUACGGUCAAUCCGCCGUUUGUUCCCA<br>CGGAGAAUCCGACGGGUUGUUACUCGCUCACAUUUAAUGUUGAUGAAAGCUGGCUACAGGAAGGC<br>CAGACGCGAAUUAUUUUUGAUGGCGUUAACUCGGCGUUUCAUCUGUGGUGCAACGGGCGCUGGG<br>UCGGUUACGGCCAGGACAGUCGUUUGCCGUCUGAAUUUGACCUGAGCGCAUUUUUACGCGCCGG<br>AGAAAACCGCCUCGCGGUGAUGGUGCUGCGUUGGAGUGACGGCAGUUAUCUGGAAGAUCAGGAU<br>AUGUGGCGGAUGAGCGGCAUUUUCCGUGACGUCUCGUUGCUGCAUAAACCGACUACACAAAUCAG<br>CGAUUUCCAUGUUGCCACUCGCUUUAAUGAUGAUUUCAGCCGCGCUGUACUGGAGGCUGAAGUU<br>CAGAUGUGCGGCGAGUUGCGUGACUACCUACGGGUAACAGUUUCUUUAUGGCAGGGUGAAACGC<br>AGGUCGCCAGCGGCACCGCGCCUUUCGGCGGUGAAAUUAUCGAUGAGCGUGGUGGUUAUGCCGA<br>UCGCGUCACACUACGUCUGAACGUCGAAAACCCGAAACUGUGGAGCGCCGAAAUCCCGAAUCUCU<br>AUCGUGCGGUGGUUGAACUGCACACCGCCGACGGCACGCUGAUUGAAGCAGAAGCCUGCGAUGU<br>CGGUUUCCGCGAGGUGCGGAUUGAAAAUGGUCUGCUGCUGCUGAACGGCAAGCCGUUGCUGAUU<br>CGAGGCGUUAACCGUCACGAGCAUCAUCCUCUGCAUGGUCAGGUCAUGGAUGAGCAGACGAUGG<br>UGCAGGAUAUCCUGCUGAUGAAGCAGAACAACUUUAACGCCGUGCGCUGUUCGCAUUAUCCGAAC<br>CAUCCGCUGUGGUACACGCUGUGCGACCGCUACGGCCUGUAUGUGGUGGAUGAAGCCAAUAUUG<br>AAACCCACGGCAUGGUGCCAAUGAAUCGUCUGACCGAUGAUCCGCGCUGGCUACCGGCGAUGAG<br>CGAACGCGUAACGCGAAUGGUGCAGCGCGAUCGUAAUCACCCGAGUGUGAUCAUCUGGUCGCUG<br>GGGAAUGAAUCAGGCCACGGCGCUAAUCACGACGCGCUGUAUCGCUGGAUCAAAUCUGUCGAUCC<br>UUCCCGCCCGGUGCAGUAUGAAGGCGGCGGAGCCGACACCACGGCCACCGAUAUUAUUUGCCCG<br>AUGUACGCGCGCGUGGAUGAAGACCAGCCCUUCCCGGCUGUGCCGAAAUGGUCCAUCAAAAAAUG<br>GCUUUCGCUACCUGGAGAGACGCGCCCGCUGAUCCUUUGCGAAUACGCCCACGCGAUGGGUAAC<br>AGUCUUGGCGGUUUCGCUAAAUACUGGCAGGCGUUUCGUCAGUAUCCCCGUUUACAGGGCGGCU<br>UCGUCUGGGACUGGGUGGAUCAGUCGCUGAUUAAAUAUGAUGAAAACGGCAACCCGUGGUCGGC<br>UUACGGCGGUGAUUUUGGCGAUACGCCGAACGAUCGCCAGUUCUGUAUGAACGGUCUGGUCUUU<br>GCCGACCGCACGCCGCAUCCAGCGCUGACGGAAGCAAAACACCAGCAGCAGUUUUUCCAGUUCCG<br>UUUAUCCGGGCAAACCAUCGAAGUGACCAGCGAAUACCUGUUCCGUCAUAGCGAUAACGAGCUCC<br>UGCACUGGAUGGUGGCGCUGGAUGGUAAGCCGCUGGCAAGCGGUGAAGUGCCUCUGGAUGUCG<br>CUCCACAAGGUAAACAGUUGAUUGAACUGCCUGAACUACCGCAGCCGGAGAGCGCCGGGCAACUC<br>UGGCUCACAGUACGCGUAGUGCAACCGAACGCGACCGCAUGGUCAGAAGCCGGGCACAUCAGCG<br>CCUGGCAGCAGUGGCGUCUGGCGGAAAACCUCAGUGUGACGCUCCCCGCCGCGUCCCACGCCAU<br>CCCGCAUCUGACCACCAGCGAAAUGGAUUUUUGCAUCGAGCUGGGUAAUAAGCGUUGGCAAUUUA<br>ACCGCCAGUCAGGCUUUCUUUCACAGAUGUGGAUUGGCGAUAAAAAACAACUGCUGACGCCGCUG<br>CGCGAUCAGUUCACCCGUGCACCGCUGGAUAACGACAUUGGCGUAAGUGAAGCGACCCGCAUUGA<br>CCCUAACGCCUGGGUCGAACGCUGGAAGGCGGCGGGCCAUUACCAGGCCGAAGCAGCGUUGUUG<br>CAGUGCACGGCAGAUACACUUGCUGAUGCGGUGCUGAUUACGACCGCUCACGCGUGGCAGCAUC<br>AGGGGAAAACCUUAUUUAUCAGCCGGAAAACCUACCGGAUUGAUGGUAGUGGUCAAAUGGCGAUU<br>ACCGUUGAUGUUGAAGUGGCGAGCGAUACACCGCAUCCGGCGCGGAUUGGCCUGAACUGCCAGC<br>UGGCGCAGGUAGCAGAGCGGGUAAACUGGCUCGGAUUAGGGCCGCAAGAAAACUAUCCCGACCG<br>CCUUACUGCCGCCUGUUUUGACCGCUGGGAUCUGCCAUUGUCAGACAUGUAUACCCCGUACGUC<br>UUCCCGAGCGAAAACGGUCUGCGCUGCGGGACGCGCGAAUUGAAUUAUGGCCCACACCAGUGGC<br>GCGGCGACUUCCAGUUCAACAUCAGCCGCUACAGUCAACAGCAACUGAUGGAAACCAGCCAUCGC<br>CAUCUGCUGCACGCGGAAGAAGGCACAUGGCUGAAUAUCGACGGUUUCCAUAUGGGGAUUGGUG<br>GCGACGACUCCUGGAGCCCGUCAGUAUCGGCGGAAUUCCAGCUGAGCGCCGGUCGCUACCAUUA<br>CCAGUUGGUCUGGUGUCAAAAAUAA |

| | |
|---|---|
| *NirFP* | CCAUCUUAGUAUAUUAGUUAAGUAUAAGAAGGAGAUAUACAUAUGGGAGAGGAUAGCGAGCUGAU<br>CUCCGAGAACAUGCACACGAAACUGUACAUGGAGGGCACCGUGAACGGCCACCACUUCAAGUGCA<br>CAUCCGAGGGCGAAGGCAAGCCCUACGAGGGCACCCAGACCUGUAAGAUCAAGGUGGUCGAGGG<br>CGGCCCUCUCCCCUUCGCCUUCGACAUCCUGGCUACCAGCUUCAUGUACGGCAGCAAAACCUUUA<br>UCAACCACACCCAGGGCAUCCCCGACUUCUUUAAGCAGUCCUUCCCUGAGGGCUUCACAUGGGAG<br>AGGAUCACCACAUACGAAGACGGGGGCGUGCUGACCGCUACCCAGGACACCAGCCUCCAGAACGG<br>CUGCCUCAUCUACAACGUCAAGAUCAACGGGGUGAACUUCCCAUCCAACGGCCCUGUGAUGCAGA<br>AGAAAACACUCGGCUGGGAGGCCAACACCGAGAUGCUGUACCCCGCUGACAGCGGUCUGAGAGG<br>CCAUAAUCAGAUGGCCCUGAAGCUCGUGGGCGGGGGCUACCUGCACUGCUCCCUCAAGACCACA<br>UACAGAUCCAAGAAACCCGCUAAGAACCUCAAGAUGCCCGGCUUCUACUUCGUGGACCGUAAACU<br>GGAAAGAAUCAAGGAGGCCGACAAAGAGACCUACGUCGAGCAGCACGAGAUGGCUGUGGCCAGGU<br>ACUGCGACCUGCCUAGCAAACUGGGGCACAGCUGA |
| *Nir*-Pep | Pepper-GUCGACGUUCACGUUGGCUUUAGUUAUUC-*NirFP* |
| 47R-cO | F30-2d×Broccoli-UCUAGAGAAUUCGGUCUCA**CAGCAGCAGCAGCAGCAGCAGCAGCAGCAGCAGC**<br>**AGCAGCAGCAGCAGCAGCAGCAGCAGCAGCAGCAGCAGCAGCAGCAGCAGCAGCAGCAGCAGC**<br>**AGCAGCAGCAGCAGCAGCAGCAGCAGCAGCAGCAGCAGCAGCAG**UUUUUUCCGCGGCUAGCA<br>UAACCCCUUGGUUCACGUUGGCUUUAGUUAUUC |
| 47R-cY | F30-2d×Broccoli-UCUAGAGAAUUCGGUCUCA**CAGCAGCAGCAGCAGCAGCAGCAGCAGCAGCAGC**<br>**AGCAGCAGCAGCAGCAGCAGCAGCAGCAGCAGCAGCAGCAGCAGCAGCAGCAGCAGCAGCAGC**<br>**AGCAGCAGCAGCAGCAGCAGCAGCAGCAGCAGCAGCAGCAGCAG**UUUUUUCCGCGGCUAGCA<br>UAACCCCUUGCGAAAAAGAGUAAAACGGCGAGGAUG |
| 47R-cN | F30-2d×Broccoli-UCUAGAGAAUUCGGUCUCA**CAGCAGCAGCAGCAGCAGCAGCAGCAGCAGCAGC**<br>**AGCAGCAGCAGCAGCAGCAGCAGCAGCAGCAGCAGCAGCAGCAGCAGCAGCAGCAGCAGCAGC**<br>**AGCAGCAGCAGCAGCAGCAGCAGCAGCAGCAGCAGCAGCAGCAG**UUUUUUCCGCGGACUAGU<br>AACUUGUUUAUUGCAGCGAAUAACUAAAGCCAACGUGAAC |
| 47R-Pep | Pepper-GAUCAUUCAUGGCAAUCUAGAGAAUUCGGUCUCAC**CAGCAGCAGCAGCAGCAGCAGCAG**<br>**CAGCAGCAGCAGCAGCAGCAGCAGCAGCAGCAGCAGCAGCAGCAGCAGCAGCAGCAGCAGCAG**<br>**CAGCAGCAGCAGCAGCAGCAGCAGCAGCAGCAGCAGCAGCAGCAGCAGCAGCAG**UUUUUUCC<br>GCGG |
